# The geometry of abstraction in hippocampus and pre-frontal cortex

**DOI:** 10.1101/408633

**Authors:** Silvia Bernardi, Marcus K. Benna, Mattia Rigotti, Jérôme Munuera, Stefano Fusi, C. Daniel Salzman

## Abstract

The curse of dimensionality plagues models of reinforcement learning and decision-making. The process of abstraction solves this by constructing abstract variables describing features shared by different specific instances, reducing dimensionality and enabling generalization in novel situations. Here we characterized neural representations in monkeys performing a task where a hidden variable described the temporal statistics of stimulus-response-outcome mappings. Abstraction was defined operationally using the generalization performance of neural decoders across task conditions not used for training. This type of generalization requires a particular geometric format of neural representations. Neural ensembles in dorsolateral pre-frontal cortex, anterior cingulate cortex and hippocampus, and in simulated neural networks, simultaneously represented multiple hidden and explicit variables in a format reflecting abstraction. Task events engaging cognitive operations modulated this format. These findings elucidate how the brain and artificial systems represent abstract variables, variables critical for generalization that in turn confers cognitive flexibility.

---

When encountering a new situation, the ability to determine right away what to think, feel, or do is a hallmark example of cognitive, emotional and behavioral flexibility. This ability relies on the fact that the world is structured. In other words, new situations in the world often share features with previously experienced ones. These shared features may correspond to hidden variables not directly observable in the environment, as well as to explicit variables. If past and future situations can be described by a small number of variables, then generalization in novel situations could be achieved by a reduced number of observations. In this scenario, the identification of few relevant variables enables generalization and can overcome the ‘curse of dimensionality’, obviating the need to enumerate and observe all possible combinations of values of all features appearing in the environment. This suggests that a process of dimensionality reduction can identify a compact set of relevant variables which exhaustively describe the environment. These variables - that correspond to features shared by multiple examples (instances) - can be represented in the brain as abstract variables, or concepts.

An account of how the brain may represent abstract variables has remained elusive, in part because we lack a precise definition of what constitutes an abstract variable. Motivated by the central role of generalization for defining abstraction, we developed analytic methods for defining when a variable is represented in an abstract format by determining the geometric properties of neural representations that support generalization. More specifically, we operationally defined a neural representation of a variable as being in an abstract format (an “abstract variable”) as a representation for which a linear neural decoder trained to report the value of the variable can generalize to new situations. These new situations are described by previously unseen combinations of the values of other variables. Within the experiment, they correspond to task conditions not used for training the linear decoder, and hence we call this ability to generalize “cross-condition generalization”.

Neural representations of an abstract variable can be obtained by representing only information related to a selected variable, and discarding all other information. However, this representation would encode only a single abstract variable. We will show that it is possible to construct representations that encode multiple variables in abstract format as defined by cross-condition generalization. To determine whether the brain represents multiple variables that are abstract according to this definition, we designed an experiment in which task conditions were described by multiple variables. Some of these variables were related to sensory inputs or motor outputs (i.e., explicit variables, such as the operant action performed and the reward outcome or value of a trial). The task also contained a “hidden” (or latent) variable defined by the temporal statistics of events and not by any particular input or output. If this hidden variable is represented in an abstract format, it would reflect a process of abstraction that involves the dimensionality reduction of sequences of events (see e.g. ^1^). Monkeys performed a serial reversal-learning task in which they switched between two un-cued contexts. Each context had distinct sets of trials containing different stimulus-response-outcome mappings (“task sets”); each set thereby could be described by a hidden variable. This task engaged a series of cognitive operations, including perceiving a stimulus, making a decision as to which action to execute, and then expecting and sensing reward to determine if an error occurred so as to update the decision process on the next trial.

Neurophysiological recordings were targeted to the hippocampus (HPC) and two parts of the pre-frontal cortex (PFC), the dorsolateral pre-frontal cortex (DLPFC) and anterior cingulate cortex (ACC). The HPC has long been implicated in generating episodic associative memories ^2–4^ that could play a central role in creating and maintaining representations of abstract variables. Indeed, studies in humans have suggested a role for HPC in the process of abstraction ^5, 6^. Neurons in ACC and DLPFC have been shown to encode rules and other cognitive information ^7–12^, but testing whether information is represented in a format that can support cross-condition generalization, especially across multiple variables, has generally not been examined (but see ^12^).

We sought to determine if during performance of this task, the targeted brain regions represent multiple variables in an abstract format. Furthermore, we examined whether the format of representations evolves as task events and concomitant cognitive operations unfold during trials. Our focus was therefore not only what information is represented by brain areas during task performance, but also on the geometric format of representations that support generalization to new conditions (an abstract format), an approach that could reveal distinctive roles of the HPC, DLPFC and ACC.

In both simulated multi-layer networks and recorded data, we analyzed the geometry of neural representations in an unbiased manner by considering all possible variables that describe equally-sized groupings of trial conditions (i.e., dichotomies of the set of all trial conditions). Neural ensembles in all 3 brain areas, and ensembles of units in simulated networks, simultaneously represented hidden and explicit variables in an abstract format (context, action and value). Notably, the representation in DLPFC of context moved into and out of an abstract format in relation to task events that engaged different cognitive operations, despite the fact that information about context remained decodable throughout. These data highlight the critical importance of characterizing the geometric format of a neural representation - not just what information is represented - in order to understand a brain region’s potential contribution to cognitive behavior relying on abstraction.

## Results

In the ensuing sections of the Results, we first present details of the behavioral task in which monkeys adjust their operant behavior in relation to changes in the hidden variable context. Next, we describe the theoretical framework and analytic methodology developed to characterize when neural ensembles represent one or more variables in an abstract format. Finally, we utilize this analytic methodology to characterize neural representations in the HPC, DLPFC, and ACC recorded during task performance, and in simulated neural networks.

### Monkeys use inference to adjust their behavior

Monkeys performed a serial-reversal learning task in which each of two blocks of trials contained four trial conditions. Three variables described each trial condition: a stimulus, and its operant and reinforcement contingencies. During the task, un-cued and simultaneous switches occurred between the blocks of trials; these switches involved changes in the contingencies of the four stimuli. Thus each block defined a context characterized by a distinct set of 4 stimulus-response-outcome mappings (task sets, with the same task sets used for all experiments). Context was thereby a hidden variable.

Correct performance for two of the stimuli in each context required releasing the button after stimulus disappearance; for the other two stimuli, the correct operant response was to continue to hold the button (Figure 1a,b; see Methods for details). For two of the stimuli, correct performance resulted in reward delivery; for the other two stimuli, correct performance did not result in reward delivery, but it did prevent both a time-out and repetition of the same unrewarded trial (Figure 1b). Without warning, randomly after 50-70 trials, the operant and reinforcement contingencies changed to those of the other context; contexts switched many times within an experiment.

**Figure 1:**
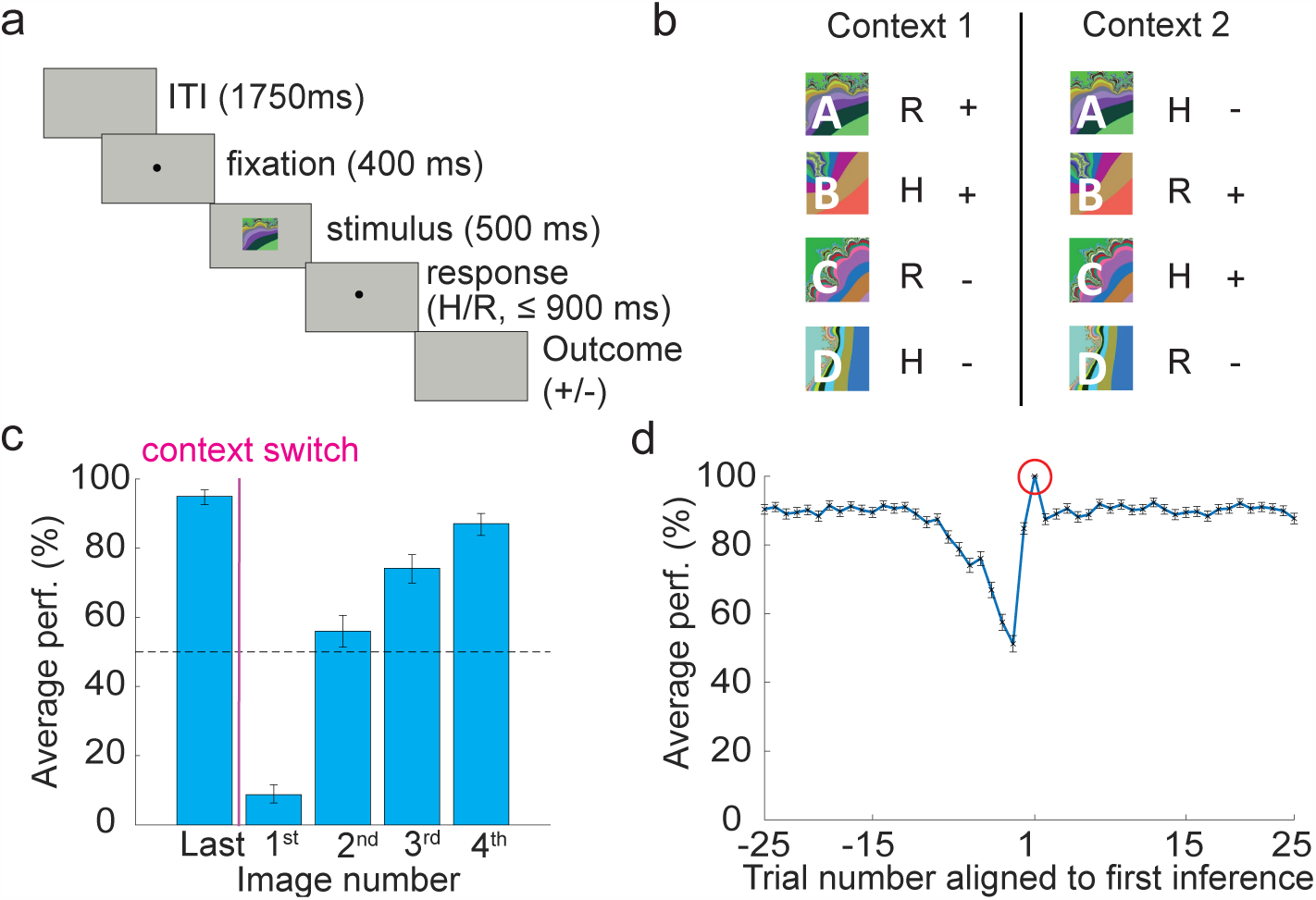
Task design and behavior. a. Sequence of events within a trial. A monkey holds down a press button, then fixates, and then views one of four familiar fractal images. A delay interval follows stimulus viewing during which the operant response (continue to hold or release the press button, respectively H and R, depending on the stimulus displayed) must be performed within 900 ms. After a trace period, a liquid reward is then delivered for correct responses for 2 of the 4 stimuli. For correct responses to the other 2 stimuli no reward is delivered but the monkey is allowed to proceed to the following trial; for incorrect responses on these trials, the trial was repeated so that monkeys could not avoid the non-rewarded trial types. b. Task scheme, stimulus response outcome map. In each of 2 contexts, 2 stimuli require the animal to continue to hold the button, while the other 2 stimuli require to release the button (H and R). Correct responses result in reward for 2 of the 4 stimuli, and no reward for the other 2 (plus or minus). Operant and reinforcement contingencies are unrelated, so neither operant action is linked to reward per se. After 50-70 trials in one context, monkeys switch to the other context and they continue to switch back-and-forth between contexts many times. c. Monkeys utilize inference to adjust their behavior. Average percent correct is plotted for the first presentation of the last image that appeared before a context switch (“Last”) and for the first instance of each image after the context switch (1-4). For image numbers 2-4, monkeys adjusted their behavior to above chance levels despite not having experienced these trials in the current context, thereby demonstrating utilization of inference. Binomial parameter estimate, bars are 95% Clopper-Pearson confidence intervals d. Average percent correct performance plotted as a function of trial number when aligning the data to the first correct trial where the monkey utilized inference (circled in red), which is defined as the first correct trial that occurs on the first presentation of either the 2nd, 3rd or 4th image type appearing after a context switch. In other words, given that Image 1 is the first image presented after a context switch, if the first presentation of image 2 after a context switch was correct, it is the first correct inference trial. If the first presentation of image 2 was incorrect, the first correct inference trial could occur on the first presentation of image 3 (if it is correct) or 4 (if it is correct, and the first presentation of both Images 2 and 3 were incorrect). Performance remains at asymptotic levels once evidence of inference is demonstrated.

Not surprisingly since context switches were un-cued, monkeys’ performance dropped to significantly below chance immediately after a context switch (see image number 1 in Fig. 1c). In principle, after this incorrect choice, monkeys could simply re-learn the correct stimulus-response associations for each image independently. Behavioral evidence indicates that this is not the case. Instead, the monkeys perform inference such that as soon as they experienced the changed contingencies for one or more stimuli upon a context switch, average performance was significantly above chance for the stimulus conditions that were not yet experienced after the context switch (see image numbers 2-4 in Fig. 1c). In this case, inference could reflect generalization to other trial types in a context after recognizing a context switch. Furthermore, as soon as monkeys exhibited evidence of inference by performing correctly on an image’s first appearance after a context switch, the monkeys’ performance was sustained at asymptotic levels (∼90% correct) for the remainder of the trials in that context. The asymptotic performance observed after a single correct inference trial indicates that monkeys subsequently performed the task as if they were in this new context (Fig. 1d). Note that it is the temporal statistics of events (trials) that defines the variable context, which is a hidden variable, not cued by any specific sensory stimulus. The only feature that the trials of Context 1 have in common is that they are frequently followed or preceded by other trials of Context 1, and the same applies to Context 2 trials. Context 1 trials rarely are followed by Context 2 trials, and vice-versa, and the monkeys behavior suggests that they exploit knowledge of these temporal statistics.

### The geometry of neural representations that encode abstract variables

Variables may be represented by a neural ensemble in many different ways. We now consider different types of representations of variables, and these types have different generalization properties. We will use these properties to define a representation of a variable in an abstract format. We first consider the hidden variable context. There exist at least two types of neural representations that encode context, but do not reflect any process of abstraction. In the first type, each neuron in an ensemble responds only to a single stimulus-response-outcome combination, i.e. to a specific trial condition, or instance of a context. In this case, the representations do not contain information about how the different instances (trial conditions) are linked together to define the two contexts. However, the neural ensemble clearly provides distinct patterns of activity for trials in each of the two contexts, and the variable context can be easily decoded. In the second type of neural representations, the firing rate of each neuron is random for each stimulus-response-outcome combination. Again, the patterns of activity corresponding to trials in each context will be distinct.

Figure 2a depicts the second type of representation in the firing rate space. In this space, each coordinate axis is the firing rate of one neuron, and hence, the total number of axes is as large as the number of recorded neurons. Each point in Figure 2a represents a vector containing the average activity of 3 neurons for each trial condition within a specified time window. The geometry of a representation is then defined by the arrangement of all the points corresponding to the different experimental conditions. In the random case, if the number of neurons is sufficiently large, the pattern of activity corresponding to each combination of stimulus-response-outcome will be unique. If trial-by-trial variability in firing rate (i.e. noise) is not too large, even a simple linear decoder can decode context. This type of random representation will allow for a form of generalization, as a decoder trained on a subset of trials can likely generalize to held-out trials. Hence, this form of generalization is not sufficient to characterize abstraction because the representations were constructed from random patterns of responses to trial conditions. Random patterns of activity for each trial condition cannot reflect the links between the different instances of the contexts (i.e. the temporal statistics of stimulus-response-outcome combinations that define context). Thus despite encoding context, this type of representation cannot be considered to represent context in an abstract format.

**Figure 2:**
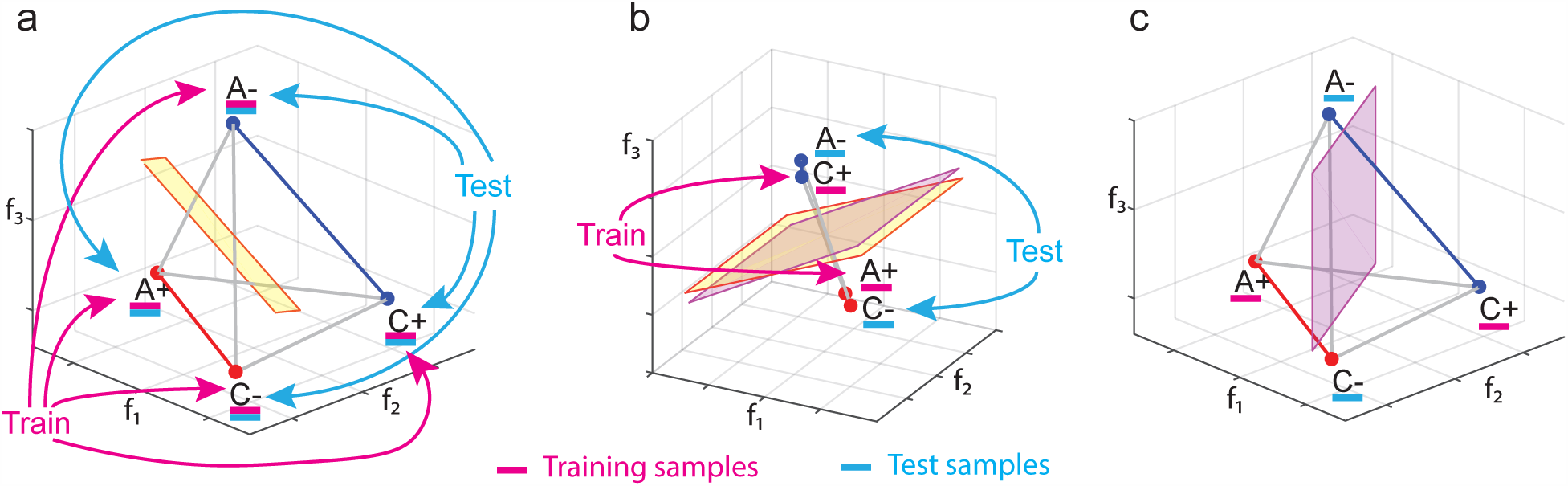
The geometry of abstraction. Each panel depicts in the firing rate space points that represent the average firing rate of a population of three neurons in one experimental condition. For simplicity we show only 4 of the 8 trial conditions. These conditions are labeled by stimulus identity (A,C) and value (+,-). a. A random representation (points are at random locations in the firing rate space), which allows for decoding of context. The yellow plane represents a linear decoder that separates the 2 points of context 1 (red) from the 2 points of context 2 (blue). The decoder is trained on all conditions (purple) and then tested on held out trials (cyan). b. Abstraction by clustering and cross condition generalization performance (CCGP). The points are clustered according to context. A linear classifier is trained to discriminate context on only two of the conditions (one from each context, in the case shown the rewarded conditions, purple). Its generalization performance is tested on the remaining conditions not used for training (here the unrewarded conditions, cyan). The resulting test performance (CCGP) will depend on the choice of training conditions. In general, the separating plane when trained on a subset of conditions (purple) is different from the one obtained when all conditions are used for training (yellow). For this clustered geometry, both planes are shown and they are very similar. c. Random representations are not abstract. CCGP is at chance level in the case of random representations, but, as shown in (a), context can be decoded by a linear classifier trained on all types of task conditions. CCGP is at chance because when the decoder is trained on the rewarded conditions, the remaining points have equal probability of being on either side of the plane. The separating plane for CCGP (purple) is very different from the separating plane of (a), obtained by training the decoder on all conditions.

To construct a neural representation of context that is in an abstract format, we need to incorporate into the geometry information about the links between different instance, in our case the temporal statistics of the stimulus-response-outcome combinations. One way to accomplish this is to cluster together patterns of activity that correspond to trials in the same context. Figure 2b illustrates this clustered geometry for 4 trial conditions. The points in the firing rate space that correspond to trials from context 1 cluster around a single point. The other points, for trials occurring in context 2, form a different cluster. Thus the patterns of activity corresponding to the two conditions that define a context are similar to each other, and are different from the patterns that define the other context.

Clustering is a geometric arrangement that permits an important and distinct form of generalization that we use to define when a neural ensemble represents a variable in an abstract format. We propose that this format supports a fundamental aspect of cognitive flexibility, the ability to generalize to novel situations. In the case of our analysis, the ability to generalize to novel conditions is analogous to being able to decode a variable in experimental conditions (stimulus-response-outcome combinations) that have not been used for training. This type of generalization is illustrated in Figure 2b. Here a linear decoder is trained to decode context on the trial conditions of the two contexts in which the animal is rewarded. Then the decoder is tested on the trials in which the animal did not receive a reward, which represents a novel situation for the decoder as it never had experienced the unrewarded trial conditions. The clustered geometric arrangements of the points ensures that the decoder successfully generalizes to the new trial conditions.

In marked contrast to clustered geometric arrangements, random responses to trial conditions do not allow for this form of generalization (Figure 2d). In the case of random responses, the two test points corresponding to the unrewarded conditions will have the same probability of being on either side of the separating plane that represents the decoder trained on other two points. Nonetheless, as we illustrated, the random representations encode context, as a linear decoder can decode it perfectly well (Figure 2c). We designate the performance of decoders in classifying variables when testing and training on different types of trial conditions as cross-condition generalization performance (CCGP) (see also ^12^). We use CCGP as a metric of the degree to which a variable is represented in an abstract format. Finally, we consider a variable represented in an abstract format when CCGP is significantly different from the one in which the points from the same trial conditions would be at random locations (see Methods).

### The geometry of multiple abstract variables

The clustering geometry allows a single variable to be encoded in an abstract format. How can neural representations encode multiple variables in an abstract format at the same time? Consider the example illustrated in Figure 3a, which depicts again a neural representation in the firing rate space. This representation was constructed by assuming that two neurons are highly specialized at encoding one variable each: Neuron 3 encodes only context, and Neuron 2 encodes only reward value. Thus the points for the trials from the two contexts lie on two parallel lines, as do the points for the trials from the two reward values. In fact, in this case a clustered geometry exists in each subspace spanned by an axis of a specialized neuron. In the different subspaces, context and value are each represented in an abstract format. This geometry allows for high CCGP for both context and value (see Figure 3a).

**Figure 3:**
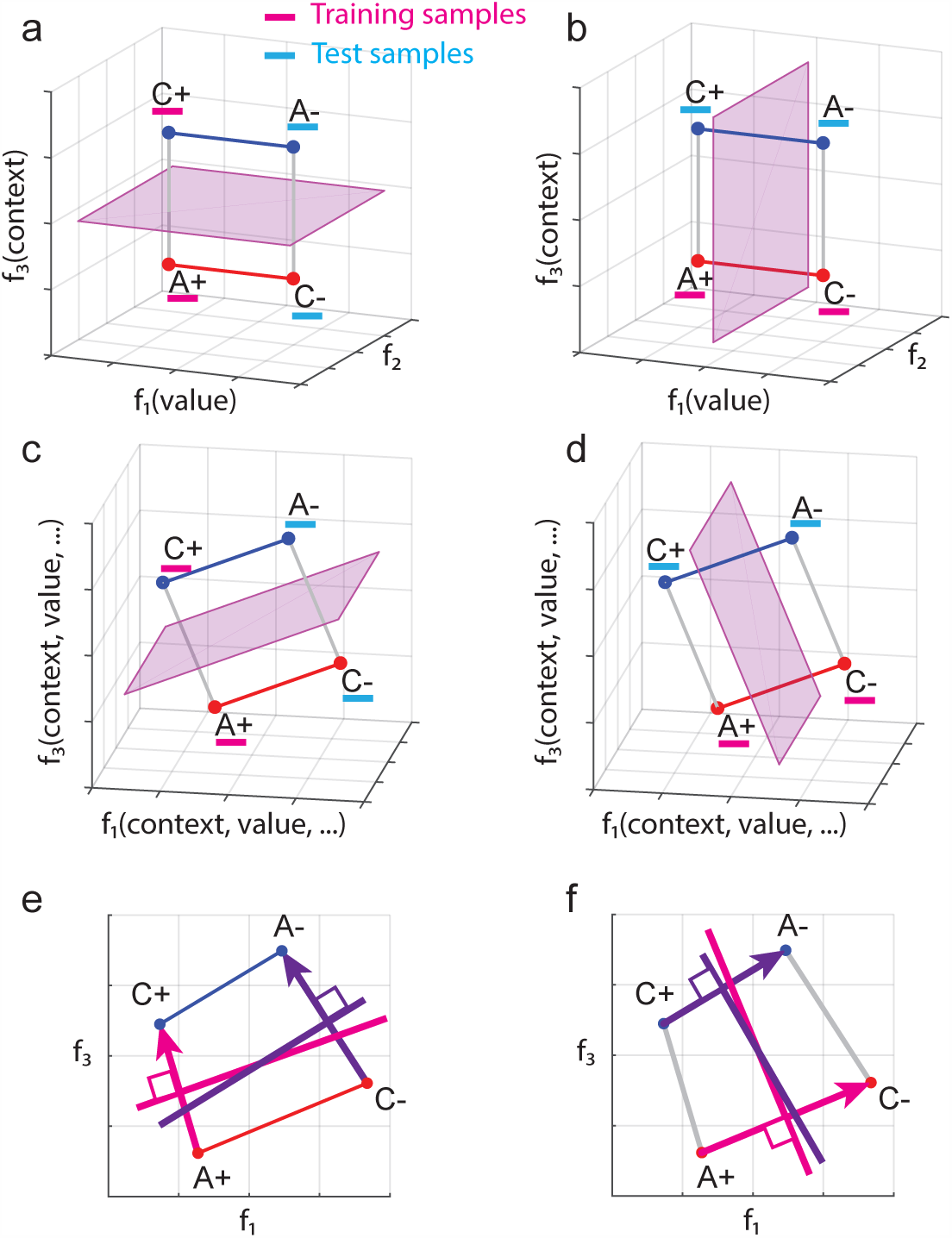
Encoding multiple abstract variables. a-f. Schematic of the firing rate space of three neurons. a,b. one neuron is specialized for encoding context (*f*_3_ axis) and one for encoding value (*f*_1_). The points of each context are in one of the two low-dimensional manifolds (lines in this case) that are parallel. These neural representations allow for cross-condition generalization for both context (a) and value (b). However, neurons that are highly specialized to encode only context are rarely observed in the data (see Suppl. Info. S4). c,d. The neural representation geometry in panel a,b is rotated in firing rate space, leading to each neuron’s exhibiting linear mixed selectivity. Even though no neurons are specialized to encode only context, in terms of decoding as well as cross-condition generalization using linear classifiers, this case is equivalent to that shown in panel a,b. e,f. Schematic explanation of the parallelism score (PS). The firing rate space has been rotated to simplify the explanation (the *f*_2_ axis is perpendicular to the page). The representations of c,d have been distorted to take into account noise in the data. e. Training a linear classifier to decode context on the two rewarded conditions leads to the magenta separating hyperplane, which is defined by a weight vector orthogonal to it. Similarly, training on the unrewarded conditions leads to the dark purple hyperplane and weight vector. If these two weight vectors are close to parallel, the corresponding classifiers are more likely to generalize to the conditions not used for training. The parallelism score (PS) is defined as the cosine of the angle between these coding vectors, maximized over all possible ways of pairing up the conditions (see Methods for details). f. Same as e but for the variable value.

In our dataset, neurons that only respond to a single variable are rarely observed (Supp Fig of all neurons), a finding consistent with many studies that have demonstrated that neurons more commonly exhibit mixed selectivity for multiple variables ^13, 14^. However, the generalization properties of the representations of context and reward value shown in Figure 3d are preserved even when we rotate the points in the firing rate space (see Figure 3e). Indeed, under the assumption that a decoder is linear, performance will not change if a linear operation like rotation is performed on the data points. Now each of the 3 neurons responds to more than one task-relevant variable, and, more specifically, exhibits linear mixed selectivity ^13, 14^ to context and reward value. Using a similar construction, it is possible to represent as many abstract variables as the number of neurons. However, additional limitations would arise from the amount of noise that might corrupt these representations.

### Measuring abstraction

The serial reversal-learning task contains 8 types of trials (stimulus-response-outcome combinations). There exist 35 different ways of dividing the 8 types of trials into two different groups of 4 trial conditions (i.e. 35 dichotomies). Each of these dichotomies corresponds to a variable that could be in an abstract format. Three of these variables are easily interpretable because they describe the context, reward value and correct action associated with each stimulus in each of the two contexts. We sought to understand which of the 35 dichotomies were encoded in the HPC, ACC, and DLPFC, and which of these 35 variables were represented in an abstract format. Variables were considered encoded if they could be decoded by a classical linear classifier, which, as usual, was trained on a subset of trials from ALL conditions and tested on held out trials. As we just reviewed, however, a variable may be encoded but not in an abstract format. To assess which variables were in an abstract format and to characterize the geometry of the recorded neural representations, we used two quantitative measures.

The first measure is CCGP, which we used to define abstract variables. CCGP can be applied to neural data by training a linear decoder to classify any dichotomy on a subset of conditions, and testing classification performance on conditions not used for training. The second measure, called the parallelism score (PS) is related to CCGP, but it focuses on specific aspects of the geometry. In particular, the PS quantifies the degree to which coding directions are parallel when training a decoder to classify a variable for different sets of conditions. Consider the case depicted in Figure 3e,f. Two different lines (which would be hyperplanes in a higher dimensional plot) are obtained when a decoder is trained to classify context using the two points on the left (the rewarded conditions, magenta) or the two points on the right (unrewarded conditions, dark purple). The two lines representing the hyperplanes are almost parallel, indicating that this geometry will allow for good cross condition generalization (high CCGP). The extent to which these lines (hyperplanes) are aligned can be quantified by calculating the coding directions (the arrows in the figure) that are orthogonal to the lines. We designate the degree to which these coding vectors are parallel as the PS. High PS usually predicts high CCGP (see Section M5).

### HPC, DLPFC, and ACC represent variables in an abstract format

We recorded the activity of 1378 individual neurons in two monkeys while they performed the serial reversal learning task. Of these, 629 cells were recorded in HPC (407 and 222 from each of the two monkeys, respectively), 335 cells were recorded in ACC (238 and 97 from each of the two monkeys), and 414 cells were recorded in DLFPC (226 and 188 from the two monkeys). Our initial analysis of neural data focused on the time epoch immediately preceding stimulus presentation. If monkeys employ a strategy in which context information is integrated with stimulus identity information to form a decision on each trial, then context information should still be stored during this time epoch. Furthermore, information about recently received rewards and performed actions may also be present (see Discussion).

In a 900 ms time epoch ending prior to the onset of a visual response on the current trial, individual neurons in all 3 brain areas exhibited mixed selectivity with respect to the task conditions on the prior trial, with diverse patterns of responses observed (see Figure S2); note that information about the current trial is not yet available during this time epoch. We took an unbiased approach to understanding which variables were represented in neural ensembles in each brain area, as well as whether the representation of any given variable defined by a specific dichotomy was in an abstract format. The traditional application of a linear neural decoder revealed that most of the 35 variables could be decoded from neural ensembles in all the brain areas, including the context, value and action of the previous trial (Fig. 4a). However, very few of these variables were represented in an abstract format at levels different from chance, as quantified by the CCGP. In fact, the variables with the highest CCGP were the variables corresponding to context and value in all 3 brain areas, as well as action in DLPFC and ACC. Action was not in an abstract format in HPC despite being decodable using traditional methods. The representation of the abstract variables did not preferentially rely on the contribution of neurons with selectivity for only one variable, indicating that neurons with linear mixed selectivity for multiple variables likely play a key role in generating representations of variables in an abstract format (see Section S4). Consistent with the CCGP analyses, the highest PSs observed in DLPFC and ACC corresponded to the 3 variables for context, value, and action, with all significantly greater than expected by chance. In HPC, the two highest PSs were for context and value, with action having a PS not significantly different than chance. The geometric architecture revealed by the CCGP and the PS analyses, can be visualized by projecting the data into a 3D space using multidimensional scaling (MDS), see Fig. S3.

**Figure 4:**
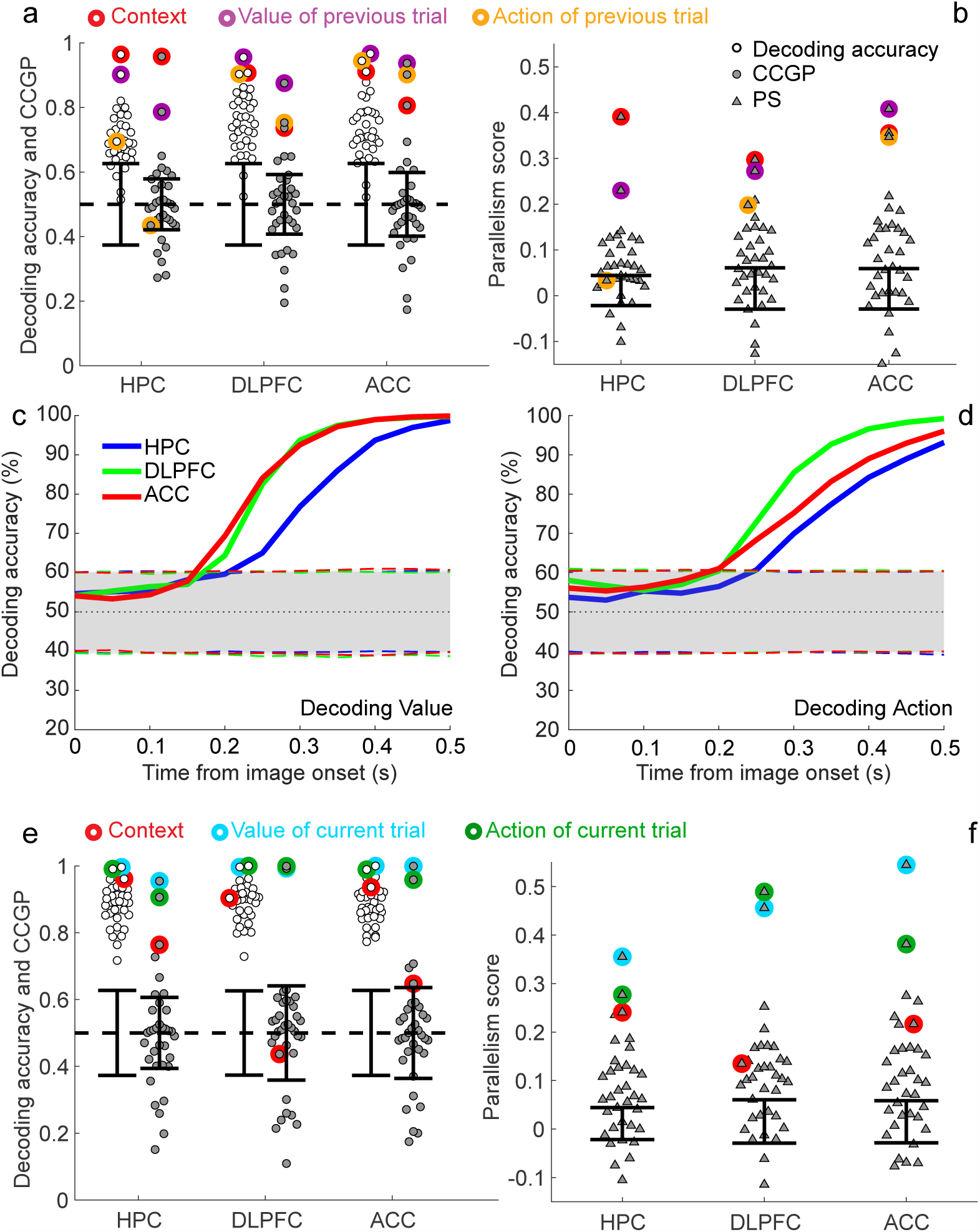
a-b. CCGP, decoding accuracy and PS for the variables that correspond to all 35 possible dichotomies shown separately for each brain area in a 900 ms time epoch beginning 800 ms before image presentation. The points corresponding to the context, value and action of the previous trial are highlighted using circles of different colors. Context and value are represented in an abstract format in all three brain areas, but action is abstract only in PFC (although it can be decoded in HPC (see also Figures S3,S4 to understand the arrangement of the points in the firing rate space). c, d. Decoding accuracy for value and action aligned on stimulus onset. e,f. CCGP, decoding accuracy and PS for all 35 dichotomies in the time interval from 100ms to 1000ms after stimulus onset. Error bars are *±* two standard deviations around chance level as obtained from a geometric random model (CCGP) or from a shuffle of the data (decoding accuracy and PS).

Notice that the hidden variable context was represented in an abstract format in all 3 brain areas just before neural responses to stimuli on the current trial occur. An analysis that only assesses whether the geometry in the firing rate space resembles clustering, similar to what has been proposed in ^5^, would lead to the incorrect conclusion that context is strongly abstract only in HPC (see Supplementary S2 and Fig. S6). The representation of context in an abstract format in ACC and DLPFC relies on a geometry revealed both by CCGP and by the PS, but missed if only quantifying the degree of clustering. By employing CCGP and PS to assess the geometry of neural representations, we also show how multiple variables can be stored in an abstract format simultaneously.

### The dynamics of the geometry of neural representations during task performance

The different task events in the serial-reversal learning task engage a series of cognitive operations, including perception of the visual stimulus, formation of a decision about whether to release the button and reward expectation. We next characterized how these task events modulated the geometry of the neural representations. Neurophysiological data indicate that the computations reflecting a decision occur rapidly after the stimulus appears in the recorded brain areas. Shortly after stimulus appearance, decoding performance for expected reinforcement outcome and the to-be-performed action on the current trial rises from chance levels to asymptotic levels in all 3 brain areas (Figure 4c,d). Decoding performance for value and action rises the most slowly in HPC, suggesting that the monkeys’ decisions are not first represented there. By contrast, in the 1 sec after stimulus appearance, decoding performance for context remained high in all 3 brain areas.

We next analyzed the geometry of neural representations during the time interval in which the planned action and expected trial outcome first become decodable. We focused on a 900 ms window beginning 100 ms after stimulus onset. We again took an unbiased approach, and considered all possible 35 dichotomies. In this time interval, nearly all of them were decodable using a traditional decoding approach (Figure 4e). When we examined the geometry of the neural representations, multiple variables were again simultaneously represented in an abstract format, and the highest CCGPs and PSs were for value and action in all 3 brain areas (Figure 4e,f). Strikingly, context was not represented in an abstract format in DLPFC; in ACC, the CCGP indicated that context was only very weakly abstract. Recall, however, that the geometry of the representation of context evolves prior to the presentation of the stimulus on the next trial, as context is in an abstract format in DLPFC and ACC during this time interval (Fig. 4a,b). In HPC, context was maintained much more strongly in an abstract format after stimulus appearance, as well as prior to stimulus appearance on the next trial. Overall, the CCGP results were largely correlated with the PS characterizing the geometry of the firing rate space. Together these findings indicate that task events both engage a series of different cognitive operations to support performance and modulate the format of neural representations. The analytic approach reveals a fundamental difference in how the format of the hidden variable context is represented in HPC compared to PFC brain areas once computations predicting a behavioral decision and expected reinforcement outcome become evident in PFC.

### Abstraction in multi-layer neural networks trained with back-propagation

We next asked whether neural representations observed in a simple neural network model trained with back-propagation have similar geometric features as those observed experimentally. We designed our simulations such that the 8 classes of inputs contained no structure that reflected a particular dichotomy, but the network had to output two arbitrarily selected variables corresponding to two specific dichotomies. We hypothesized that forcing the network to output these two variables would break the symmetry between all dichotomies. Similar neural representations would then be generated for inputs that share the same output, leading to abstract representations of the output variables. This simulated network would therefore provide us with a way of generating abstract representations of selected variables which we could use to benchmark our analytic methods. In particular, our methods should identify only the output variables as being represented in an abstract format.

We trained a two layer network using back-propagation (see Figure 5a) to read an input representing a handwritten digit between 1 and 8 (from the MNIST dataset) and to output whether the input digit is odd or even, and, at the same time, whether the input digit is large (*>* 4) or small (≤4) (Figure 5b). Parity and magnitude are the two variables that we hypothesized could be abstract. We tested whether the learning process would lead to high CCGP and PS for parity and magnitude in neural representations in the last hidden layer of the network. If these variables are in an abstract format, then the abstraction process would be similar to the one studied in the experiment in the sense that it involves aggregating together inputs that are visually dissimilar (e.g. the digits ‘1’ and ‘3’, or ‘2’ and ‘4’). Analogously, in the experiment very different sequences of events (visual stimulus, operant action and value) are grouped together into what we defined as contexts, sets of stimulus-response-outcome contingencies that constitute a hidden variable.

**Figure 5:**
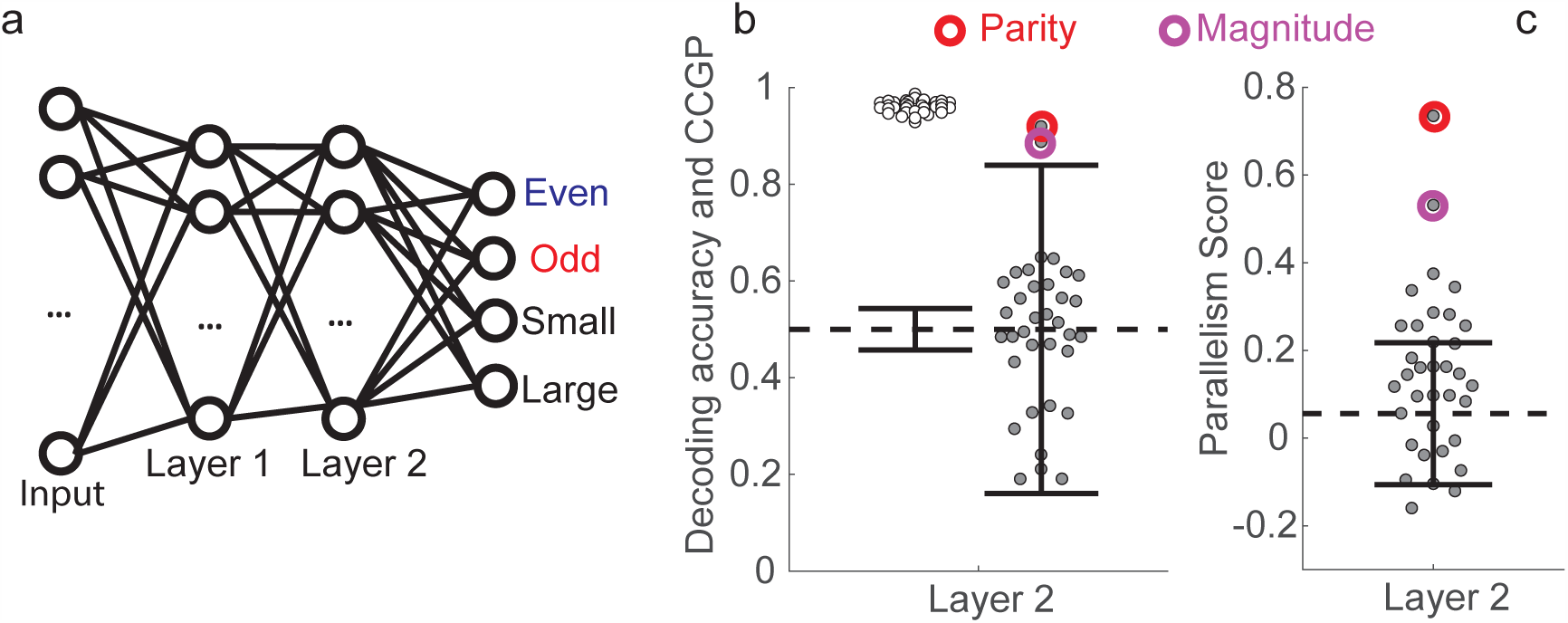
Simulations of a multi-layer neural network reveal that the geometry of the observed neural representations can be obtained with a simple model. a. Diagram of the network architecture. The input layer receives gray-scale images of MNIST handwritten digits with 784 pixels. The two hidden layers have 100 units each, and in the final layer there are two pairs of output units corresponding to two binary variables. The network is trained using back-propagation to simultaneously classify inputs (we only use the images of digits 1-8) according to whether they depict even/odd and large/small digits. b. Cross-condition generalization performance and decoding accuracy for the variables corresponding to all 35 possible balanced dichotomies when the second hidden layer is read out. Only the two dichotomies corresponding to parity and magnitude are significantly different from a geometric random model (chance level: 0.5; the two solid black lines indicate plus/minus two standard deviations). The decoding performance is high for all dichotomies, and hence inadequate to identify the variables stored in an abstract format. c. Same as b, but for the parallelism score (PS), with error bars (plus/minus 2 standard deviations) obtained from a shuffle of the data. Both the CCGP and PS allow us to identify the abstract variables actually used to train the network. See Fig. S5 for a multi-dimensional scaling visualization of the neural representations.

Just as in the experiments, we computed both the CCGP and the PS for all possible dichotomies of the eight digits. In Figure 5d,e we show the decoding accuracy, the CCGP and the PS for all dichotomies. The largest CCGP and PS correspond to the parity dichotomy, and the second largest values correspond to the magnitude dichotomy. For these two dichotomies, both the CCGP and the PS are significantly different from those of the random models. No other dichotomies exhibit a CCGP value that is statistically significant, despite the fact that all dichotomies can be decoded using a traditional decoding approach. This analysis shows that the CCGP and the PS identify the abstract variables that correspond to the dichotomies encoded in the output. Note that the geometry of the representations in the last layer would actually allow the network to perform classification of any dichotomy, as the decoding accuracy is close to 1 for every dichotomy. Thus abstraction is not necessary for the network to perform tasks that require outputs corresponding to any of the 35 dichotomies. However, the simulations show that only parity and magnitude are represented in an abstract format.

Next, we asked whether a neural network trained to perform a simulated version of our experimental task would also exhibit a similar geometry. This exercise offers the advantage of mimicking our task more closely. We used a reinforcement learning algorithm (Deep Q-Learning) to train the network. This technique uses a deep neural network representation of the state-action value function of an agent trained with a combination of temporal-difference learning and back-propagation refined and popularized by ^15^. The use of a neural network is ideally suited for a comparative study between the neural representations of the model and the recorded neural data. As commonly observed in Deep Q-learning, neural representations displayed significant variability across runs of the learning procedure. However, in a considerable fraction of runs, the neural representations during a modeled time interval preceding stimulus presentation recapitulate the main geometric features that we observed in the experiment (see Suppl. Info. S6). In particular, after learning, the hidden variable context is represented as an abstract variable in the last layer, despite not being explicitly represented in the input, nor in the output. Moreover, the neural representations of the simulated network encode multiple task-relevant variables in an abstract format simultaneously, consistent with the observation that hidden and explicit variables are represented in an abstract format in the corresponding time interval in the experiment.

## Discussion

Neuroscientists have often focused on what information is encoded in a particular brain area without considering in what format the information is stored. However, cognitive and emotional flexibility relies on our ability to represent general concepts in a format that enables generalization upon encountering new situations. Here we developed two analytic approaches for characterizing the geometric format of neural representations to understand how variables may be represented in an abstract format to support generalization.

Our data reveal that neural ensembles in DLPFC, ACC and HPC represent multiple variables in an abstract format simultaneously during performance of a serial-reversal learning task. This finding held for the action planned and executed to reach a decision, the reinforcement expected or recently received, and for the hidden variable context. The content and format of representations of variables changed as task events engaged the cognitive operations needed to perform the task. In the time period just prior to stimulus onset, context and the reward outcome of the previous trial were in abstract format in all 3 brain areas. Despite being decodable in all brain areas, the action of the previous trial was not in an abstract format in HPC, unlike in ACC and DLPFC. After an image appeared, neural representations of the planned action and the expected reward occur more rapidly in DLPFC and ACC than in HPC, suggesting that these pre-frontal areas may play a more prominent role in the decision process. Notably, value and action were in abstract format in all brain areas shortly after image onset, but context was not abstract in the DLPFC despite being decodable. In ACC, context was only weakly abstract despite being strongly decodable. Two implementations of simulated multi-layer networks exhibited a similar geometry of representations of abstract variables. These results highlight how the format in which a variable is represented can distinguish between the coding properties of brain areas, even when the content of information is present in those areas.

### The potential role of neural representations of action and value of previous trial

Although information about the value and action of the previous trial are represented in all three brain areas right before stimulus presentation on the current trial, if the animal did not make a mistake on the previous trial, this information is not needed to perform the current trial correctly. However, at the context switch, the reward received on the previous trial is the only feedback from the external world that indicates that the context has changed. Therefore, reward value is essential when adjustments in behavior are required, suggesting that storing the recently received reward could be beneficial. Consistent with this notion, we show in simulations that value becomes progressively more abstract as the frequency of context switches increases (see S6 and Figure S19). In addition, monkeys occasionally make mistakes that are not due to a context change. To discriminate between these occasional errors and those due to a context change, information about value is not sufficient and information about the previously performed action could be essential for deciding correctly the operant response on the next trial. Conceivably, the abstract representations of reward and action may also afford the animal more flexibility in learning and performing other tasks. Previous work has shown that information about recent events is represented whether it is task-relevant or not (see e.g. ^16, 17^). This storage of information (a memory trace) may even degrade performance on a working memory task ^18^, but presumably the memory trace might be beneficial in other scenarios that demand cognitive flexibility in a range of real-world situations.

### Dimensionality and abstraction in neural representations

Dimensionality reduction is widely employed in many machine learning applications and data analyses because it leads to better generalization. In our theoretical framework, we constructed representations of abstract variables that are indeed relatively low-dimensional, as the individual neurons exhibit linear mixed selectivity ^13, 14^. These constructed representations have a dimensionality that is equal to the number of abstract variables that are simultaneously encoded. Consistent with this, the neural representations recorded in the time interval preceding the presentation of the stimulus are relatively low-dimensional (Supplementary S5).

Previous studies showed that the dimensionality of neural representations can be maximal (monkey PFC ^13^), very high (rodent visual cortex^19^), or as high as it can be given the structure of the task ^20^. These results seem to be inconsistent with what we now report. However, dimensionality is not a static property of neural representations; in different epochs of a trial, dimensionality can vary significantly (see e.g. ^21^). Dimensionality has been observed to be maximal in a time interval in which all the task-relevant variables had to be mixed non-linearly to support task performance ^13^. Here we analyzed two time intervals. In the first time interval, which preceded visual responses to images, the variables that are encoded do not need to be mixed. However, during a later time interval beginning 100 ms after stimulus presentation, the dimensionality of the neural representations increases significantly (Supplementary S5), suggesting that the context and the current stimulus are mixed non-linearly later in the trial, similar to prior observations ^13, 14, 19, 22, 23^. This may account for why context is not abstract in DLPFC, as mixing of context and stimulus identity information could underlie the computations required for decision formation on this task. We note that there is not necessarily a strong negative correlation between dimensionality and the degree of abstraction. Distortions of the idealized abstract geometry can significantly increase dimensionality, providing representations that preserve some ability to generalize, but at the same time support operations requiring higher dimensional representations (see Supplementary Information S8).

### The role of abstraction in reinforcement learning

Abstraction is an active area of research in Reinforcement Learning (RL), as it provides a solution for the notorious “curse of dimensionality”; that is, the exponential growth of the solution space required to encode all states of the environment ^24^. Most abstraction techniques in RL can be divided into two main categories: ‘temporal abstraction’ and ‘state abstraction’. Temporal abstraction is the workhorse of Hierarchical Reinforcement Learning ^25–27^ and is based on the notion of temporally extended actions (or options). Temporal abstraction can thereby be thought of as an attempt to reduce the dimensionality of the space of action sequences: instead of having to compose policies in terms of long sequences of actions, the agent can select options that automatically extend for several time steps.

State abstraction methods are most closely related to our work. In brief, state abstraction hides or removes information about the environment not critical for maximizing the reward function. Typical instantiations of this technique involve information hiding, clustering of states, and other forms of domain aggregation and reduction ^28^. Our use of neural networks as function approximators to represent a decision policy constitutes a state abstraction method (see Results and Supplementary Information S6). Here the inductive bias of neural networks induces generalization across inputs sharing a feature, allowing them to mitigate the curse of dimensionality. The modeling demonstrates that neural networks induce the type of abstract representations observed in our data. Furthermore, the modeling suggests that our analysis techniques could be useful to elucidate the geometric properties underlying the success of Deep Q-learning neural networks trained to play 49 different Atari video games with super-human performance ^15^.

### Other forms of abstraction in the computational literature

The principles that we have delineated for representing variables in abstract format are reminiscent of recent work in computational linguistics. This work has suggested that difficult lexical semantic tasks can be solved by word embeddings, which are vector representations whose geometric properties reflect the meaning of linguistic tokens ^29, 30^. Recent forms of word embeddings exhibit linear compositionality that makes the solution of analogy relationships possible via linear algebra ^29, 30^. For example, shallow neural networks that are trained in an unsupervised way on a large corpus of documents end up organizing the vector representations of common words such that the difference of the vectors representing ‘king’ and ‘queen’ is the same as the difference of the vectors for ‘man’ and ‘woman’^29^. These word embeddings, which can be translated along parallel directions to consistently change one feature (e.g. gender, as in the previous example), clearly share common coding principles with the abstract representations that we describe. This type of vector representation predicts fMRI BOLD signals measured while subjects are presented with semantically meaningful stimuli ^31^.

A different approach to extracting compositional features in an unsupervised way relies on variational Bayesian inference to learn to infer interpretable factorized representations of some inputs ^1, 32–35^. These methods can disentangle independent factors of variations of a variety of real-world datasets, and it will be interesting to apply our analytical methods to gain insight into the functioning of these algorithms.

The generation of neural representations of variables in an abstract format is also critical for many other sensory, cognitive and emotional functions. For example, in vision, the creation of neural representations of objects that are invariant with respect to their position, size and orientation in the visual field is a typical abstraction process that has been studied in machine learning applications (see e.g. ^36, 37^) and in brain areas involved in representing visual stimuli (see e.g. ^38, 39^). This form of abstraction is sometimes referred to as ‘untangling’ ^40, 41^ because the retinal representations of objects correspond to manifolds with a relatively low intrinsic dimensionality but are highly curved and tangled together and then become “untangled” in visual cortex. Untangling typically requires a series of transformations that either increase the dimensionality of the representations by projecting into a higher dimensional space, or decrease their dimensionality by extracting relevant features. Abstraction as we defined it also implies an analogous dimensionality reduction.

One important difference between our work and studies on abstraction in vision (e.g. ^40, 41^) is that often in vision, the final representation is required to be linearly separable for only one specific classification task (e.g. car vs non-car). The geometry of the representation of “nuisance” variables that describe the features of the visual input not relevant for the task is not studied systematically, unlike in this paper. The nuisance variables typically are not under control when a complex visual object is placed in a natural scene, and these variables are often difficult to describe. On the contrary, the variables in our experiment that we analyzed are simple binary variables, allowing us to study systematically all possible dichotomies. Recent studies have investigated neural representations in the visual system using a measure similar to our CCGP ^42, 43^.

## Conclusions

Our study delineates an approach for distinguishing between neural ensembles that represent information, and those ensembles that also represent information in an abstract format that enables cross-condition generalization. This distinction affords the ability to ascribe differential functions to brain areas based on the format of neural representations, and not merely on the information they represent. Traditionally a variable is considered to be encoded in a neural ensemble if the variable modulates the neural activity. Here we showed the advantages of going beyond this concept of encoding by considering the geometry of neural representations. This requires knowing not only how a variable modulates neural activity, but also how other variables affect neural representations and their geometry. Future studies must focus on the specific neural mechanisms that account for the formation and modulation of neural representations of variables in abstract format in the context of a broader range of tasks, which is fundamentally important for any form of learning, for executive functioning, and for cognitive and emotional flexibility.

## Supporting information

MDS plot for ACC

MDS plot for DLPFC

MDS plot for HPC

## Acknowledgements

We are grateful to L.F. Abbott and R. Axel for many useful comments on the manuscript. This project is supported by the Simons Foundation, and by NIMH (1K08MH115365, R01MH082017). SF and MKB are also supported by the Gatsby Charitable Foundation, the Swartz Foundation, the Kavli foundation and the NSF’s NeuroNex program award DBI-1707398. JM is supported by the Fyssen Foundation. SB received support from NIMH (1K08MH115365, T32MH015144 and R25MH086466), and from the American Psychiatric Association and Brain & Behavior Research Foundation young investigator fellowships.

## Methods

### M1 Task and Behavior

Two male rhesus monkeys (Macaca mulatta; two males, 8 and 13 kg respectively) were used in these experiments. All experimental procedures were in accordance with the National Institutes of Health guide for the care and use of laboratory animals and the Animal Care and Use Committees at New York State Psychiatric Institute and Columbia University. Monkeys performed a serial-reversal learning task in which they were presented one of four visual stimuli (fractal patterns). Stimuli were consistent across contexts and sessions, presented in random order. Each trial began with the animal holding down a button and fixating for 400 ms (Fig. 1a). If those conditions were satisfied, one of the four stimuli was displayed on a screen for 500 ms. In each context, correct performance for two of the stimuli required releasing the button within 900 ms of stimulus disappearance; for the other two, the correct operant action was to continue to hold the button. For 2 of the 4 stimuli, correct performance resulted in reward delivery; for the other 2, correct performance did not result in reward. If the monkey performed the correct action, a trace interval of 500 ms ensued followed by the liquid reward or by a new trial in the case of non-rewarded stimuli. If the monkey made a mistake, a 500 ms time out was followed by the repetition of the same trial type if the stimulus was a non-rewarded one. In the case of incorrect responses to rewarded stimuli, the time-out was not followed by trial repetition and the monkey simply lost his reward. After a random number of trials between 50 and 70, the context switched without warning. Upon a context switch, operant contingencies switched for all images, but for two stimuli the reinforcement contingencies did not change, in order to maintain orthogonality between operant and reinforcement contingencies. A different colored frame (red or blue) for each context appears on the edges of the monitor on 10 percent of the trials, randomly selected, and only on specific stimulus types (stimulus C for context 1 and stimulus D for context 2). This frame never appeared in the first five trials following a contextual switch. All trials with a contextual frame were excluded from all analyses presented.

### M2 Electrophysiological Recordings

Recordings began only after the monkeys were fully proficient in the task and performance was stable. Recordings were conducted with multi-contact vertical arrays electrodes (v-probes, Plexon Inc., Dallas, TX) with 16 contacts spaced at 100 *µ*m intervals in ACC and DLPFC, and 24 contacts in HPC, using the Omniplex system (Plexon Inc.). In each session, we individually advanced the arrays into the three brain areas using a motorized multi-electrode drive (NAN Instruments). Analog signals were amplified, band-pass filtered (250 Hz - 8 kHz), and digitized (40 kHz) using a Plexon Omniplex system (Plexon, Inc.). Single units were isolated offline using Plexon Offline Sorter (Plexon, Inc.). To address the possibility that overlapping neural activity was recorded on adjacent contacts, or that two different clusters visible on PCA belonged to the same neuron, we compared the zero-shift cross-correlation in the spike trains with a 0.2 ms bin width of each neuron identified in the same area in the same session. If 10 percent of spikes co-occurred, the clusters were considered duplicated and one was eliminated. If 1-10 percent of spikes co-occurred, the cluster was flagged and isolation was checked for a possible third contaminant cell. Recording sites in DLPFC were located in Brodmann areas 8, 9 and 46. Recording sites in ACC were in the ventral bank of the ACC sulcus (area 24c). HPC recordings were largely in the anterior third, spanning across CA1-CA2-CA3 and DG.

### M3 Selection of trials/neurons, and decoding analysis

The neural population decoding algorithm was based on a linear classifier (see e.g. ^12^) trained on pseudo-simultaneous population response vectors composed of the spike counts of the recorded neurons within specified time bins and in specific trials ^44^. The trials used in the decoding analysis are only those in which the animal responded correctly (both for the current trial and the directly preceding one), in which no context frame was shown (neither during the current nor the preceding trial), and which occurred at least five trials after the most recent context switch. We retain all neurons for which we have recorded at least 15 trials satisfying these requirements for each of the eight experimental conditions (i.e., combinations of context, value and action). Every decoding analysis is averaged across many repetitions to estimate trial-to-trial variability (as explained more in detail below). For every repetition, we randomly split off five trials per condition from among all selected trials to serve as our test set, and used the remaining trials (at least ten per condition) as our training set. For every neuron and every time bin, we normalized the distribution of spike counts across all trials in all conditions with means and standard deviations computed on the trials in the training set. Specifically, given an experimental condition *c* (i.e., a combination of context, value and action) in a time bin *t* under consideration, we generated the pseudo-simultaneous population response vectors by sampling, for every neuron *i*, the z-scored spike count in a randomly selected trial in condition *c*, which we indicate by 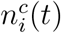 This resulted in a single-trial population response vector 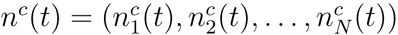, where *N* corresponds to the number of recorded neurons in an area under consideration. This single-trial response vector can be thought of as a noisy measurement of an underlying mean firing rate vector 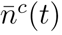, such that 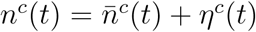, with *η*^*c*^(*t*) indicating a noise vector modeling the trial-to-trial variability of spike counts. Assuming that the trial-to-trial noise is centered at zero, we estimate the mean firing rate vectors taking the sample average: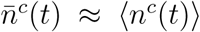, where the angular brackets indicate averaging across trials. We then either trained maximum margin (SVM) linear classifiers on the estimated mean firing rate vectors for the eight conditions in the training set (this is the approach adopted to train the decoders used to compute the CCGP, see below), or we trained such classifiers on the single-trial population response vectors generated from the training set of trials (this is what we used in all the figures that report the “decoding accuracy”). In the latter case, in order to obtain a number of trials that is large compared to the number of neurons, we re-sampled the noise by randomly picking noisy firing rates (i.e, spike counts) from among all the training trials of a given experimental condition for each neuron independently. Specifically, in this case, we re-sampled 10,000 trials per condition from the training set. While this neglected correlations between different neurons within conditions, we only had little information about these correlations in the first place, since only a relatively small numbers of neurons were recorded simultaneously. Regardless of whether we trained on estimated mean firing rate vectors or on re-sampled single-trial population response vectors, the decoding performance was measured in a cross-validated manner on 1,000 re-sampled single-trial population response vectors generated from the test set of trials. For every decoding analysis training and testing were then repeated 1,000 times over different random partitions of the trials into training and test trials. The decoding accuracies that we report were computed as the average results across repetitions.

Statistical significance of the decoding accuracy was assessed using a permutation test for classification^45^. Specifically, we repeated the same procedure just described, but at the beginning of every repetition of a decoding analysis, trials were shuffled, i.e., associated to a random condition. This is a way of estimating the probability that the population decoders that we used would have given the same results that we obtained by chance, i.e., when applied on data that contain no information regarding the experimental conditions.

In Fig. S1 we show the cross-validated decoding accuracy as a function of time throughout the trial (for a sliding 500 ms time window) for maximum margin classifiers trained only on the mean neural activities for each condition. Figs. 4a and e show similar results for linear classifiers trained on the mean firing rates in the neural data within time windows from −800 ms to 100 ms and from 100 ms to 1000 ms relative to stimulus onset, respectively.

For all analyses, data were combined across monkeys, because all key features of the data set were consistent across the two monkeys.

### M4 The cross-condition generalization performance (CCGP)

The hallmark feature of neural representations of variables in abstract format (“abstract variables”) is their ability to support generalization. When several abstract (in our case binary) variables are encoded simultaneously, generalization must be possible for all the abstract variables. We quantify a strong form of generalization using a measure we call the cross-condition generalization performance (CCGP.) We use this measure as a quantitative definition of the degree to which a variable is represented in abstract format. CCGP is distinct from traditional cross-validated decoding performance commonly employed to determine if a neural ensemble represents a variable. In traditional cross-validated decoding, the data is split up randomly such that trials from all conditions will be present in both the training and test sets. For CCGP, trials are split instead according to their condition labels, such that the training set consists entirely of trials from one group of conditions, while the test set consists only of trials from a disjoint group of conditions (see the scheme in Figure M1). In computing CCGP, we train a linear classifier for a certain dichotomy that discriminates the conditions in the training set according to some label (one of the variables), and then ask whether this discrimination generalizes to the test set by measuring the classification performance on the data from entirely different conditions, i.e. conditions not used for training the decoder. CCGP provides a continuous measure which is basically a degree of abstraction.

Given our experimental design with eight different conditions (distinguished by context, value and action of the previous trial), we can investigate variables corresponding to different balanced (four versus four condition) dichotomies, and choose one, two or three conditions from each side of a dichotomy to form our training set. We use the remaining conditions (three, two or one from either side, respectively) for testing, with larger training sets typically leading to better generalization performance. For different choices of training conditions we will in general obtain different values of the classification performance on the test conditions, and we define CCGP as its average over all possible sets of training conditions (of a given size). In Figs. 4a and e we show the CCGP (on the held out fourth condition) when training on three conditions from either side of the 35 balanced dichotomies (with dichotomies corresponding to context, value and action highlighted).

We emphasize that in order to achieve high CCGP, it is not sufficient to merely generalize over the noise associated with trial-to-trial fluctuations of the neural activity around the mean firing rates corresponding to individual conditions. Instead, the classifier has to generalize also across different conditions on the same side of a dichotomy, i.e., across those conditions that belong to the same category according to the variable under consideration.

For the CCGP analysis, the selection of trials used is the same as for the decoding analysis, except that here we retain all neurons that have at least ten trials for each experimental condition that meet our selection criteria (since the split into training and test sets is determined by the labels of the eight conditions themselves, so that for a training condition we don’t need to hold out additional test trials). We pre-process the data by z-scoring each neuron’s spike count distribution separately. Again, we can either train a maximum margin linear classifier only on the cluster centers, or on the full training set with trial-to-trial fluctuations (noise), in which case we re-sample 10,000 trials per condition, with Figs. 4a and e showing results using the latter method.

For all analyses, data were combined across monkeys, because all key features of the data set were consistent across the two monkeys.

### M5 The parallelism score (PS)

We developed a measure based on angles of coding directions to characterize the geometry of neural representations of variables in the firing rate space. Consider a pair of conditions, one from each side of a dichotomy, such as the two conditions that correspond to the presentation of stimulus A in the two contexts (here context is the dichotomy under consideration). A linear classifier trained on this pair of conditions defines a separating hyperplane. The weight vector orthogonal to this hyperplane aligns with the vector connecting the two points that correspond to the mean firing rates for the two training conditions if we assume isotropic noise around both of them. This corresponds to the coding direction for the variable under consideration (context in the example). Other coding directions for the same variable can be obtained by choosing a different pair of training conditions (e.g. the conditions that correspond to the presentation of stimulus B in the two context). The separating hyperplane associated with one pair of training conditions is more likely to correctly generalize to another pair of conditions if the associated coding directions are parallel (as illustrated in Fig. 3e,f). The parallelism score (PS) that we developed directly quantifies the alignment of these coding directions.

**Figure M1:**
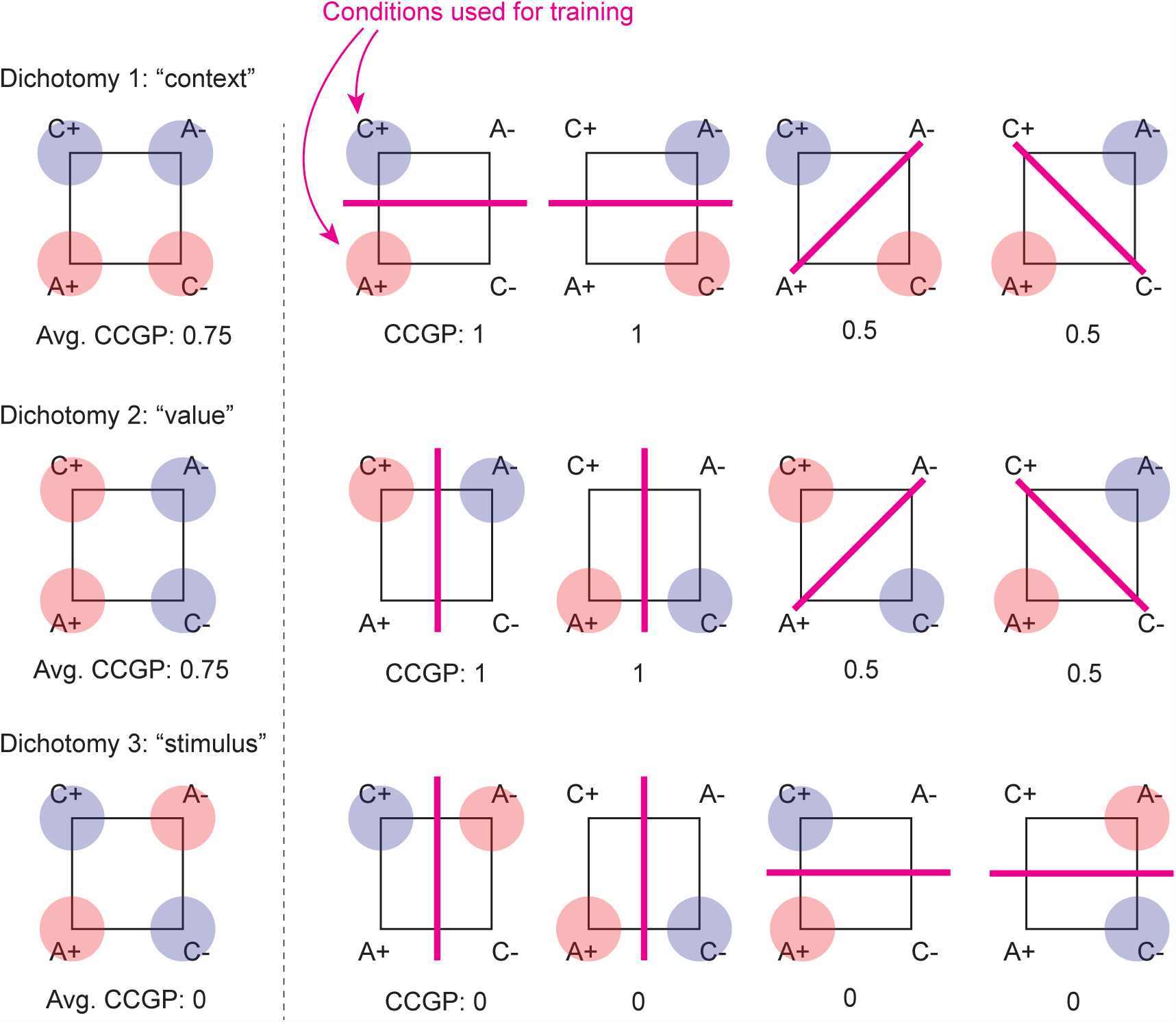
Scheme that illustrates how CCGP is computed for all dichotomies. The example has the same geometry as the representation in Figure 3a (value and context are simultaneously abstract). Each dichotomy corresponds to a different way of dividing the 4 conditions in two groups of two (different colored clouds for each group). In this example, all possible dichotomies correspond to variables that have a name (“context”, “value”, “stimulus”). However, this is not necessarily true in other situations. For each dichotomy, CCGP is computed by training a linear decoder on a subset of conditions. These are the only conditions highlighted (by shaded colored circles) in the figure. The decoder is then tested on the remaining (non-shaded) conditions. For each dichotomy there are 4 possible ways of choosing 2 conditions, one from each side of the dichotomy, as illustrated in the figure in the row next to each dichotomy. The final CCGP is obtained by computing the average test performance over all the different ways of choosing two training conditions. For this geometry, CCGP is 0.75 for the dichotomies context and value, and 0 for the dichotomy stimulus. These values are obtained under the assumption that the noise is isotropic and not too large.

If we had only four conditions as shown in Fig. 3e, there would be only two coding directions for a given variable (from the two pairs of training conditions), and we would simply calculate the cosine of the angle between them (i.e., the normalized overlap of the two weight vectors). In our experiments, there were 8 conditions whose mean firing rates we denote by *f* (*c*), with *c* = 1, 2,…, 8. A balanced dichotomy corresponds to splitting up these eight conditions into two disjoint groups of four, corresponding to the conditions to be classified as positive and negative, respectively, e.g. *G*_*pos*_ = [1, 2, 4, 7] versus *G*_*neg*_ = [3, 5, 6, 8]. To compute four unit coding vectors 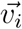 for *i* = 1, 2, 3, 4 (corresponding to four pairs of potential training conditions) for the variable associated with this dichotomy, we have to match each condition in the positive group with a unique condition in the negative group (without repetitions). We parametrize these pairings by considering all possible permutations of the condition indices in the negative group. For a particular choice of such a permutation *𝒫* the resulting set of coding vectors is given by

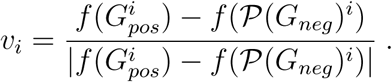

Note that we have defined the *v*_*i*_ as normalized coding vectors, since we want our parallelism score to depend only on their direction (but not on the magnitude of the un-normalized coding vectors). To compute the parallelism score from these unit coding vectors, we consider the cosines of the angles between any two of them 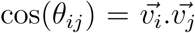 and we average these cosines over all six of these angles (corresponding to all possible choices of two different coding vectors).

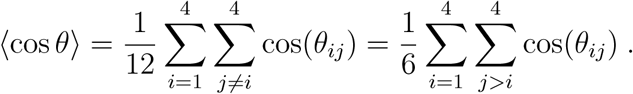

In general there are multiple ways of pairing up conditions corresponding to the two values of the variable under consideration. We don’t want to assume a priori that we know the ‘correct’ way of pairing up conditions. For example, it is not obvious that the two conditions corresponding to the same stimuli in two contexts are those that maximize the cosine between the coding vectors. It could be that cosine is larger when the conditions corresponding to a certain value are paired. In other words, to perform the analysis in an unbiased way, we should ignore the labels of the variables that define the conditions within each dichotomy (in the case of context we should just consider all conditions corresponding to context 1 and pair them in all possible ways to all conditions in context 2). So we consider all possible ways of matching up the conditions on the two sides of the dichotomy one-to-one, corresponding to all possible permutations 𝒫, and then define the PS as the maximum of the average cosine across all possible pairings/permutations. There are two such pairings in the case of four conditions, and 24 for eight conditions. In general there are (*m/*2)! pairings for *m* conditions, so if *m* was large there would be a combinatorial explosion in the obvious generalization of this definition to arbitrary *m*, which would also require averaging the cosines of (*m/*2)(*m/*2 − 1)*/*2 angles.

The parallelism score for a given balanced dichotomy in our case of eight conditions is defined as

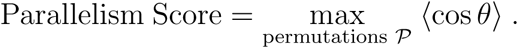

Note that this quantity depends only on the normalized coding directions (for the best possible pairing of conditions), which are simply the unit vectors pointing from one cluster center (mean firing rate for a given condition) towards another. Therefore, finding the PS doesn’t require training any classifiers, which makes it a very simple, fast computation (unless *m* is large). However, because it depends only on the locations of the cluster centers, the parallelism score ignores the shape of the noise (within condition trial-to-trial fluctuations).

The parallelism scores of all 35 dichotomies in our data (with the context, value and action dichotomies highlighted) are plotted in Figs. 4b and f. The selection of trials used in this analysis is the same as for the decoding and cross-condition generalization analyses, retaining all neurons that have at least ten trials for each experimental condition that meet our selection criteria, and z-scoring each neuron’s spike count distribution individually.

Note that a high parallelism score for one variable/dichotomy doesn’t necessarily imply high cross-condition generalization. Even if the coding vectors for a given variable are approximately parallel, the test conditions might be much closer together than the training conditions. In this case generalization would likely be poor.

In addition, for the simple example of only four conditions the orthogonal dichotomy would have a low parallelism score in such a situation (corresponding to a trapezoidal geometry).

We also emphasize that high parallelism scores for multiple variables do not guarantee good generalization (large CCGP) of one dichotomy across another one. When training a linear classifier on noisy data, the shape of the noise clouds could skew the weight vector of a maximum margin classifier away from the vector connecting the cluster centers of the training conditions. Moreover, even if this is not the case (e.g. if the noise is isotropic), generalization might still fail because of a lack of orthogonality of the coding directions for different variables. (For example, the four conditions might be arranged at the corners of a parallelogram instead of a rectangle, or in the shape of a parallelepiped instead of a cuboid for eight conditions).

In summary, while the parallelism score is not equivalent to CCGP, high scores for a number of dichotomies with orthogonal labels characterize a family of (approximately factorizable) geometries that can lead to good generalization properties if the noise is sufficiently well behaved (consider e.g. the case of the principal axes of the noise distributions being aligned with the coding vectors). For the simple case of isotropic noise, if the coding directions for different variables are approximately orthogonal to each other, CCGP will also be high.

For all analyses, data were combined across monkeys, because all key features of the data set were consistent across the two monkeys.

### M6 Random models

In order to assess the statistical significance of the above analyses we need to compare our results (for the decoding performance, abstraction index, cross-condition generalization performance, and parallelism score, which we collectively refer to as scores here) to the distribution of values expected from an appropriately defined random control model. There are various sensible choices for such random models, each corresponding to a somewhat different null hypothesis we might want to reject. We consider two different classes of random models, and for each of our abstraction analyses we choose the more conservative one of the two to compute error bars around the chance levels of the scores (i.e., we only show the one that leads to the larger standard deviation).

#### Shuffle of the data

One simple random model we consider is a shuffle of the data, in which we assign a new, random condition label to each trial for each neuron independently (in a manner that preserves the total number of trials for each condition). In other words, we randomly permute the condition labels (with values from 1 to 8) across all trials, and repeat this procedure separately for every neuron. When re-sampling artificial, noisy trials, we shuffle first, and then re-sample in a manner that respects the new, random condition labels as described above. This procedure destroys almost all structure in the data, except the marginal distributions of the firing rates of individual neurons. The error bars around chance level for the decoding performance in Figs. S1 and 4, and for the parallelism score in Figs. 4 and 5 are based on this shuffle control (showing plus/minus two standard deviations). These chance levels and error bars around them are estimated by performing the exact same decoding/PS analyses detailed in the preceding sections on the shuffled data, and repeating the whole shuffle analysis a sufficient number of times to obtain good estimates of the means and standard deviations of the resulting distributions of decoding performances/parallelism scores (e.g. for the PS we perform 1,000 shuffles).

#### Geometric random model

Another class of control models is more explicitly related to neural representations described by random geometries, and can be used to rule out a different type of null hypothesis. For the analyses that depend only on the cluster centers of the eight conditions (i.e., their mean firing rates, as e.g. for the PS), we can construct a random geometry by moving the cluster of points that correspond to different conditions to new random locations that are sampled from an isotropic Gaussian distribution. We then rescale all the vectors to keep the total variance across all conditions (the signal variance, or variance of the centroids of the clusters). Such a random arrangement of the mean firing rates (cluster centers) is a very useful control to compare against, since such geometries do not constitute abstract neural representations, but nevertheless typically allow relevant variables to be decoded (see also Figure 2). For analyses that depend also on the structure of the within condition trial-to-trial fluctuations (in particular, CCGP and decoding with re-sampled trials), our random model in addition requires some assumptions about the noise distributions. We could simply choose identical isotropic noise distributions around each cluster center, but training a linear classifier on trials sampled from such a model would essentially be equivalent to training a maximum margin classifier only on the cluster centers themselves. Instead, we choose to preserve some of the noise structure of the data by moving the (re-sampled) noise clouds to the new random position of the corresponding cluster and performing a discrete rotation around it by permuting the axes separately for each condition. We basically shuffled the neuron labels in a different way for each condition. This shuffling corresponds to a discrete rotation of each noise cloud (i.e. the cloud of points that represents the set of all trials for one specific condition). The rotations are random and independent for each condition. While the structure of the signal is completely destroyed by generating a random set of cluster centers for the eight conditions, the within condition noise structure (but not the correlations across conditions) is retained in these models. If our scores are significantly different from those obtained using this random model, we can reject the null hypothesis that the data were generated by sampling a random isotropic geometry with the same total signal variance (i.e. the variance across the different cloud centers) and similarly shaped noise clouds as in the data. The error bars around chance level for the CCGP in Figs. 4 and 5 are derived from this geometric random control model by constructing many such random geometries and estimating the standard deviation of the resulting CCGPs.

#### Random models and the analysis of different dichotomies

One might be tempted to consider the scores for the 35 different dichotomies as a set of score values that defines a random model. Indeed, this could be described as a type of permutation of the condition labels, but only between groups of trials belonging to the same condition (thus preserving the eight groups of trials corresponding to separate conditions), as opposed to the random permutation across all trials performed in the shuffle detailed above. However, there are clearly correlations between the scores of different dichotomies (e.g. because the labels may be partially overlapping, i.e., not orthogonal). Therefore, we should not think of the set of scores for different dichotomies as resulting from a random model used to assess the probability of obtaining certain scores from less structured data. After all, the different dichotomies are simply different binary functions to be computed on the same neural representations, without changing any of their essential geometric properties. Instead, the set of scores for the 35 dichotomies allows us to make statements about the relative magnitude of the scores compared to those of other variables that may also be decodable from the data and possibly abstract, as shown in the form of bee-swarm plots in Figs. 4 and 5.

### M7 Simulations of the multi-layer network

The two hidden layer network depicted in Figure 5 contains 768 neurons in the input layer, 100 in each hidden layer and four neurons in the output layer. We used eight digits (1-8) of the full MNIST data set to match the number of conditions we considered in the analysis of the experiment. The training set contained 48128 images and the test set contained 8011 digits. The network was trained to output the parity and the magnitude of each digit and to report it using four output units: one for odd, one for even, one for small (i.e., a digit smaller than 5) and one for large (a digit larger than 4). We trained the network using the back-propagation algorithm `‘train’` of matlab (with the neural networks package). We used a tan-sigmoidal transfer function (`‘tansig’` in matlab), the mean squared normalized error (`‘mse’`) as the cost function, and the maximum number of training epochs was set to 400. After training, we performed the analysis of the neural representations using the same analytical tools that we used for the experimental data, except that we did not z-score the neural activities, since they were simultaneously observed in the simulations.

The description of the methods to model our task using a reinforcement learning algorithm (Deep Q-learning) appears below in Suppl. Info. S6.

## Supplementary Information

### S1 Supplementary figures

**Figure S1:**
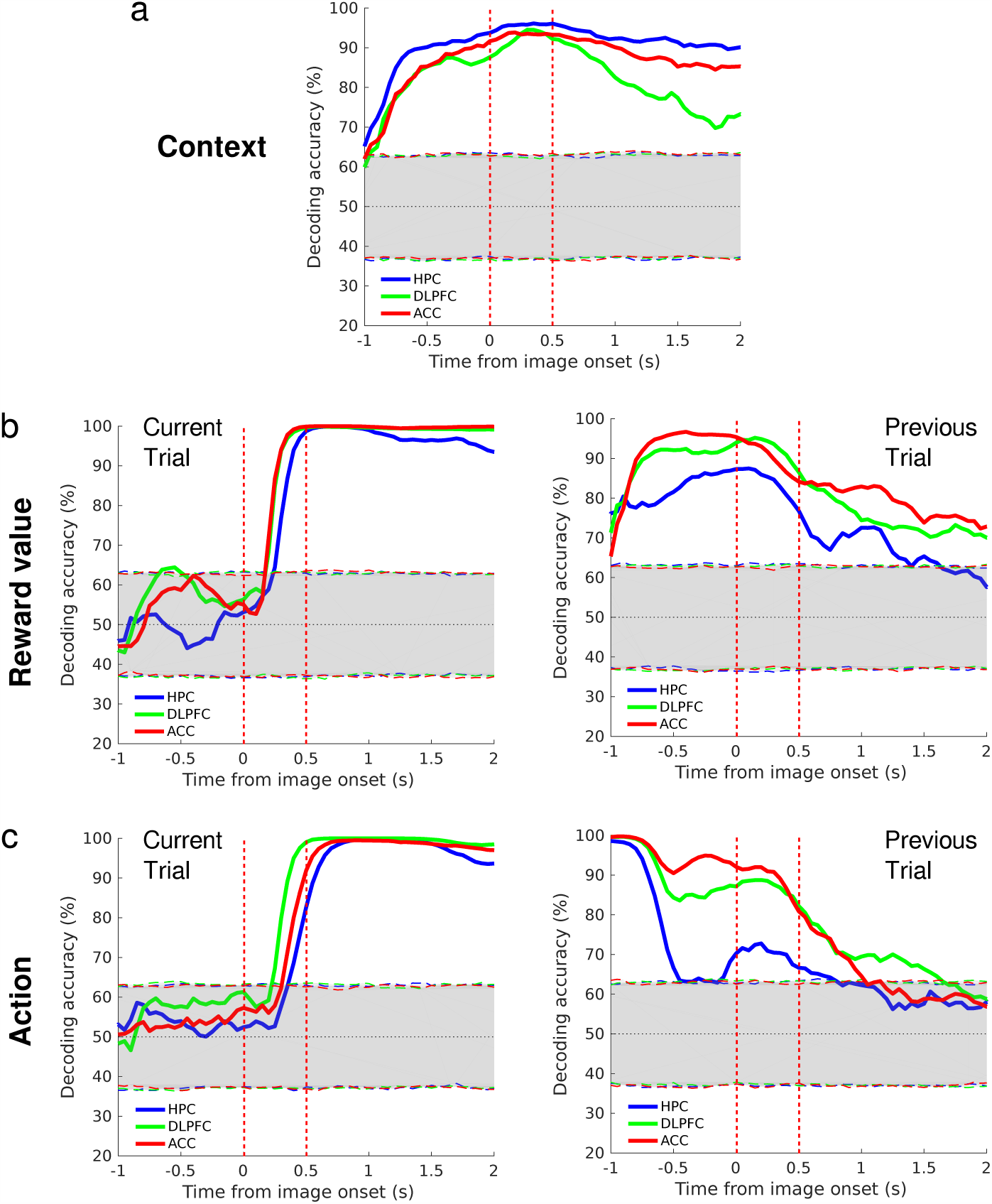
Population level encoding of task-related variables. a-e. Performance of a linear decoder plotted a function of time relative to image onset for classifying a task-relevant variable. a. Context on the current trial. b. Reinforcement outcome on current (left) and prior (right) trials. c. Operant action on current (left) and prior (right) trials. The decoding performance was computed in a 500-ms sliding window stepped every 50 ms across the trial for the three brain areas separately (blue, HPC; red, ACC; green, DLPFC). Dashed lines indicate two standard deviations above and below the mean of the distribution of accuracies obtained with a permutation test done by shuffling trials 1000 times (bootstrap). The image is displayed on the screen from time 0 to 0.5 sec. Analyses were run only on correct trials at least 5 trials after a context switch.

**Figure S2:**
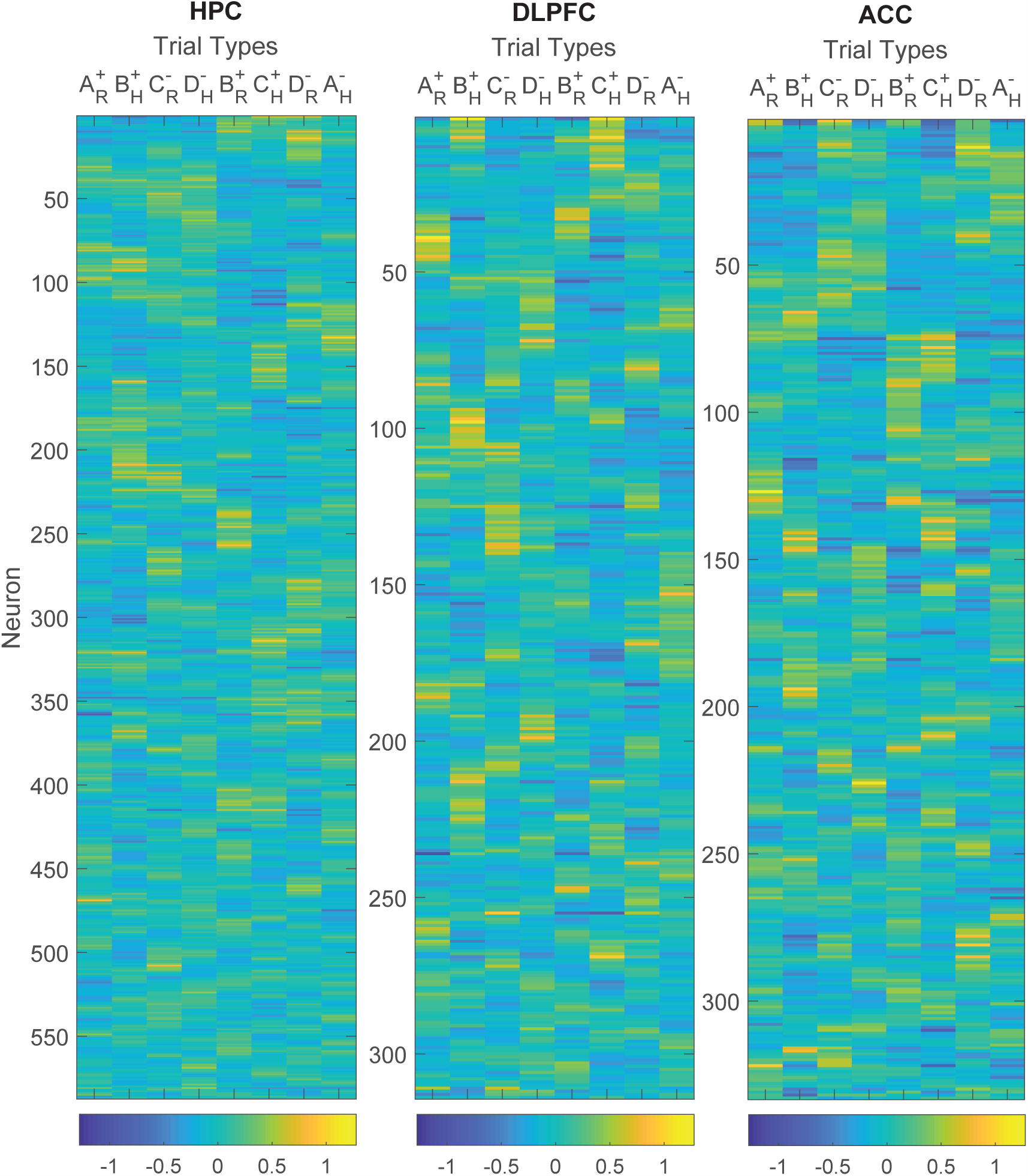
The activity of all neurons recorded for at least ten trials per condition after excluding: 1) those trials performed incorrectly; 2) those trials in which a contextual frame was shown either for the current or the previous trial; and 3) those trials that occurred less than five trials after a context switch). Z-scored firing rates are calculated in the 900 ms time window that starts 800ms before stimulus onset. Each row represents the activity of an individual neuron. Different columns correspond to different trial conditions (i.e., the stimulus-action-outcome sequence preceding the interval). Neurons are ordered such that for adjacent rows, the eight-dimensional vectors of z-scored activities point in similar directions. The responses are very diverse in all three brain areas.

**Figure S3:**
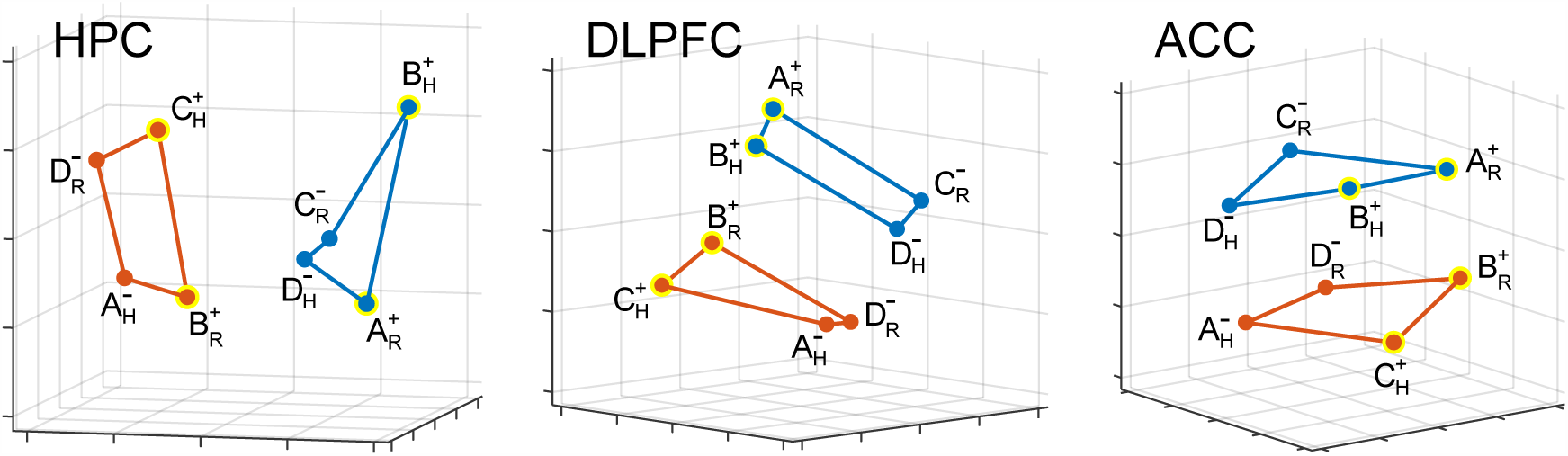
The geometry of the recorded neural representations. Multi-dimensional scaling plots (using Euclidean distances on z-scored spike count data in the 900 ms time window that starts 800ms before stimulus onset) showing the dimensionality-reduced firing rates for different experimental conditions in the three brain areas recorded from: HPC, DLPFC and ACC. The labels refer to value (+*/*−) and operant action (R/H) corresponding to the previous trial. Yellow rings are rewarded conditions. While there is a fairly clean separation between the two context sub-spaces, other variables are encoded as well and the representations are not strongly clustered. Note that the context sub-spaces appear to be approximately two-dimensional (i.e., of lower dimensionality than expected for four points in random positions). See also the videos in the Supplementary Material.

**Figure S4:**
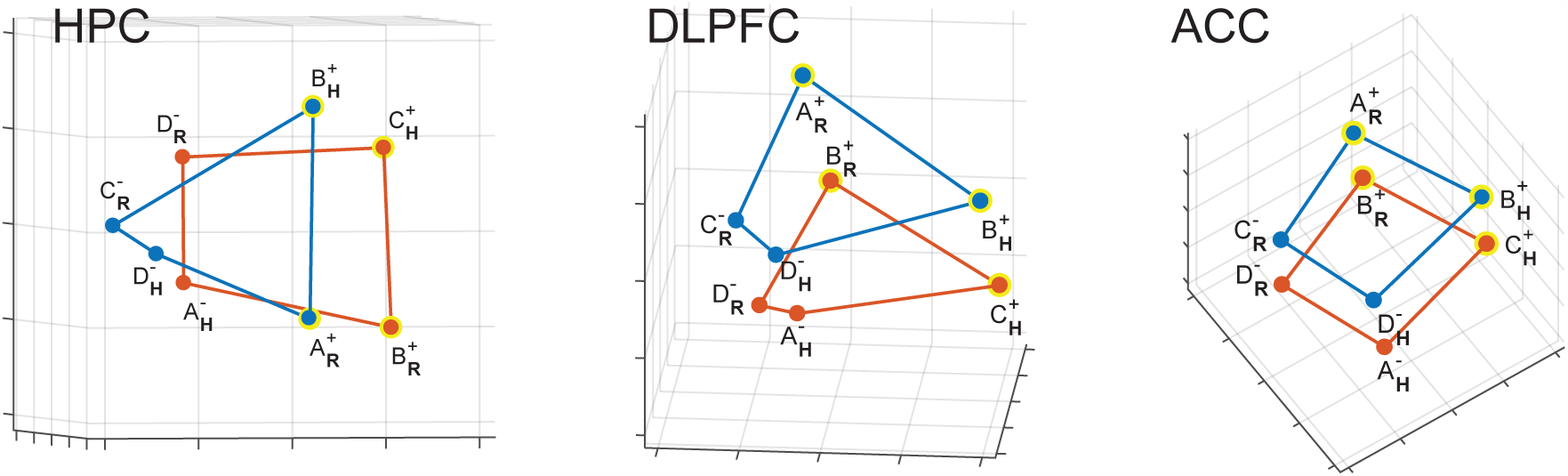
Representation of action of the previous trial in the three brain areas. The MDS plots of Figure S3 are rotated to highlight the dependence on the action of the previous trial (H=Hold, R=Release; in boldface). In DLPFC and ACC the points corresponding to the conditions with the same action are on the same side of the plot. In these areas, action is decodable and in an abstract format. In HPC all the 8 points are distinct, and action is decodable. However, it is not in an abstract format. Indeed, the H and R points are along two almost orthogonal diagonals. Notice that value (+/−symbols) and context (red/blue; less clear from this perspective) are in an abstract format in all three brain areas.

**Figure S5:**
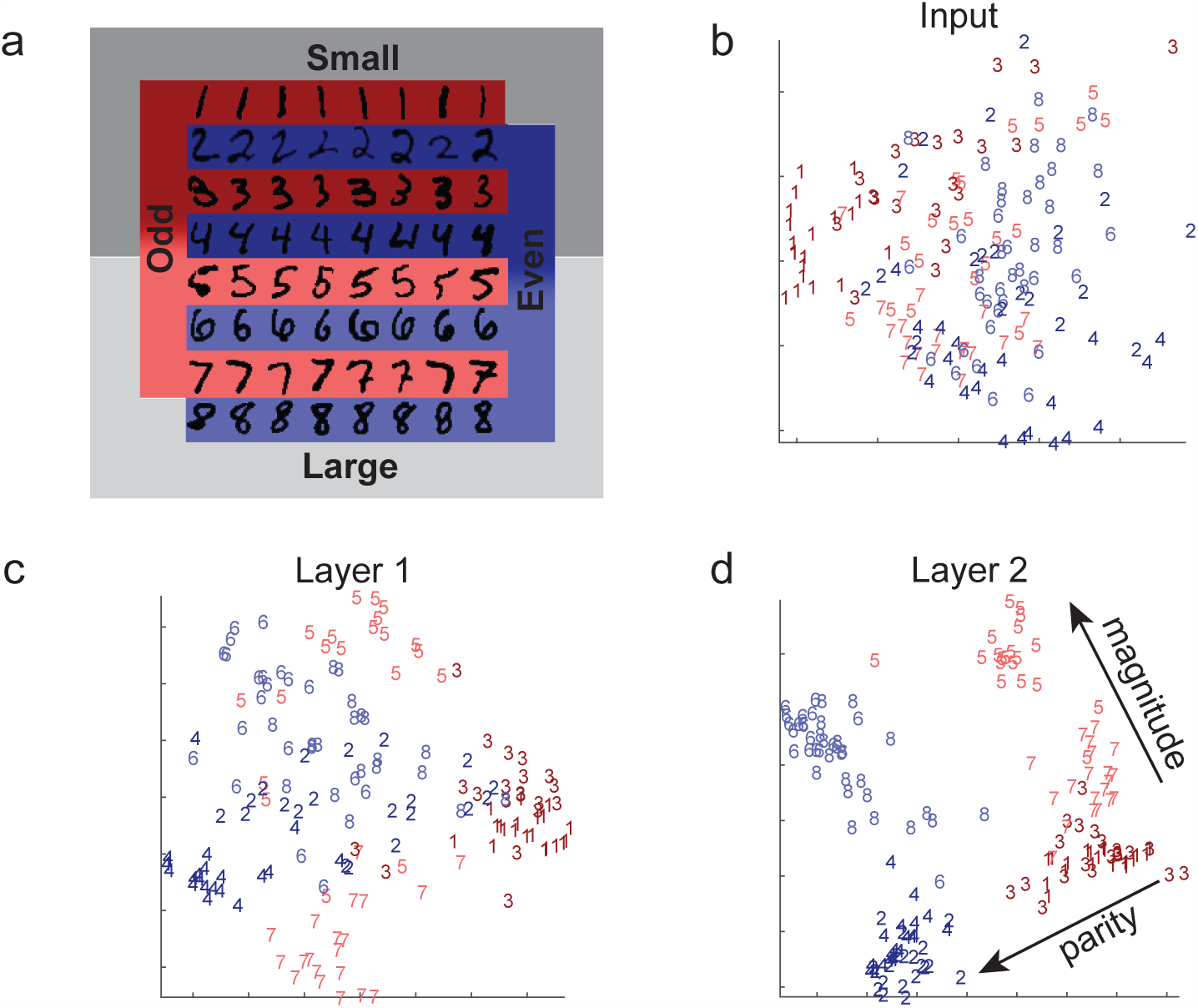
Simulations of a multi-layer neural network reveal that the geometry of the observed neural representations can be obtained with a simple model. a. Schematic of the two discrimination tasks using the MNIST dataset. The network is trained using back-propagation to simultaneously classify inputs (we only use the images of digits 1-8) according to whether they depict even/odd and large/small digits. The colors indicate parity, and shading indicates the magnitude of the digits (darker for smaller ones). b-d. Two-dimensional MDS plots of the representations of a subset of images in the input (pixel) space (b), as well as in the first (c) and second hidden layers (d). In the input layer there is no structure apart from the accidental similarities between the pixel images of certain digits (e.g. ones and sevens). However, in the first, and even more so in the second, layer, a clear separation between digits of different parities and magnitudes emerges in a geometry with consistent and approximately orthogonal coding directions for the two variables. This suggests a simultaneously abstract representation for both variables.

#### Multidimensional scaling (MDS)

MDS plots (Figs. S3, S4, S5b,c,d, and S6a) are obtained as follows. Within a chosen time bin the activity of each neuron across all conditions is z-scored. It is then averaged across trials within each condition to obtain the firing rate patterns for each condition. These patterns are then used to construct an *n*_*c*_*×n*_*c*_ dissimilarity matrix (where *n*_*c*_ is the number of conditions), which simply tabulates the Euclidean distance between firing rate patterns for each pair of conditions. This dissimilarity matrix is then centered, diagonalized, and projected along the first 3 eigenvectors rescaled by the squared root of the corresponding eigenvalues, in accordance with the Classical Multidimensional Scaling algorithm ^46^.

### S2 Frequently asked questions about our measures of abstraction

#### Can we use a clustering index to measure abstraction?

A simple way to achieve an abstract neural representation would be to cluster together the activity patterns corresponding to the conditions on either side of a certain dichotomy (defining a binary variable). For example, if the spike count patterns in a certain brain area were equal for all four stimuli in context one, and similarly coincided (at a different value) for context two, this area would exhibit an abstract representation of context, at the expense of not encoding any information about stimulus identity, operant action or reward value.

We can assess the degree of clustering by comparing the average distance between the mean neural activity patterns corresponding to the conditions on the same side of a dichotomy (within or intra-group) to the average distance between those on opposite sides of the dichotomy (between or inter-group). For balanced (four versus four) dichotomies of the eight experimental conditions (context, value and action of the previous trial), there are 16 inter-group distances, and 12 intragroup distances (six on each side of the dichotomy) that contribute. We define the ratio of the average between group distance to the average within group distance as the **abstraction index**, which measures *the degree of clustering of a set of neural activity patterns associated with a certain dichotomy*. In the absence of any particular geometric structure (such as clustering), we would expect these two average distances to be equal, resulting in an abstraction index of one, while clustering would lead to values larger than one.

Fig. S6 shows the abstraction index for the context variable computed from the measured neural activity patterns. As above for decoding, we retain only correct trials without a contextual frame that didn’t occur within 5 trials of a context switch for this analysis. We z-score the overall activity distribution (across all task conditions) of each neuron before computing the mean activity pattern of each condition, and use a simple Euclidean distance metric (we verified that employing a Mahalanobis distance metric instead of the Euclidean distance metric yields similar results).

Clustering can also identify an abstract variable (as verified by CCGP) when the geometry of a neural representation differs from that shown in Figure 2b. For example, when the points of 4 different conditions are arranged in a square, as in Figure 3, the inter-class distances are slightly larger than the intra-class distances (the diagonals connecting opposite points are larger than the length of the side of the square). The geometry is thereby clearly less clustered than in the case discussed in Figure 2b, but nonetheless the analysis could lead to the correct conclusion that the representation is abstract. However, the abstraction index, though larger than 1, would be significantly smaller than in the case of clustering illustrated in Figure 2b, and we do not know how robust this measure would be to noise.

**Figure S6:**
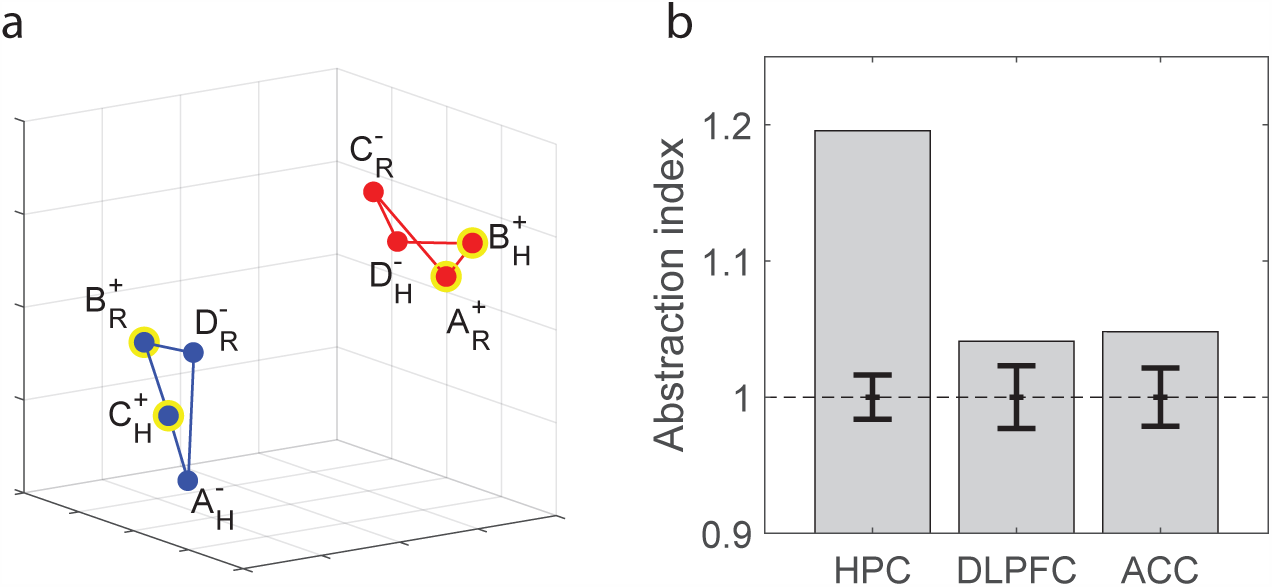
Abstraction by clustering. a) Schematic of firing rate space in the case of clustering abstraction. Due to the clustering, the average within-context distance is shorter than the mean between-context distance. b) Abstraction index for the context dichotomy (ratio of average between-context distance to average within-context distance using a simple Euclidean metric) for the z-scored neural firing rates recorded from HPC, DLPFC and ACC, averaged over a time window of −800ms to 100ms relative to stimulus onset. The error bars are plus/minus two standard deviations around chance level (unit abstraction index), obtained from a shuffle of the data.

We emphasize that although clustering can identify variables in an abstract format, there are several situations in which a clustering analysis fails to identify abstract variables, even though these variables are identified by CCGP. For example, in the geometry in Figure S7, a cube is squeezed along the vertical direction. Here context is represented in an abstract format because CCGP would be close to 1 when a decoder is trained on 6 conditions, 3 per context, and tested on the remaining ones, so long as noise is not too large. However, an analysis of clustering would not identify context as abstract. This is because when the vertical side of the rectangular cuboid is *d* and the sides of the squares are 1, then for any *d <* 0.634 the average intra-group distance is larger than the average inter-group distance. In these cases, the abstraction index is less than 1. The geometry depicted in Figure S7 is not an unlikely situation because different abstract variables are typically encoded with different strengths (in the referenced figure, action and value are represented more strongly than context).

#### How are our measures of abstraction such as Cross-condition Generalization Performance related to generalization performance in classification taks?

We motivate our CCGP measure of abstraction operationally: abstraction of one variable is quantified in terms of performance at predicting the value of the variable in novel conditions. In particular, we measure CCGP of the neural response vector representation of context (or any other variable corresponding to a dichotomy) by training a weak classifier (a linear one) to decode context from response vectors in a subset of training experimental conditions, and then testing the trained classifier on a held out subset of experimental conditions (which is analogous to “novel” conditions). Intuitively, the accuracy of the classifier in predicting context on the held out experimental conditions quantifies how “consistently” context is being encoded across conditions. So, for instance CCGP would be very low for response patterns in generic random position. Indeed, it would be straight-forward to train a linear classifier with high accuracy (in a low noise regime), but the same classifier would perform at chance level on the response patterns recorded on held out conditions.

**Figure S7:**
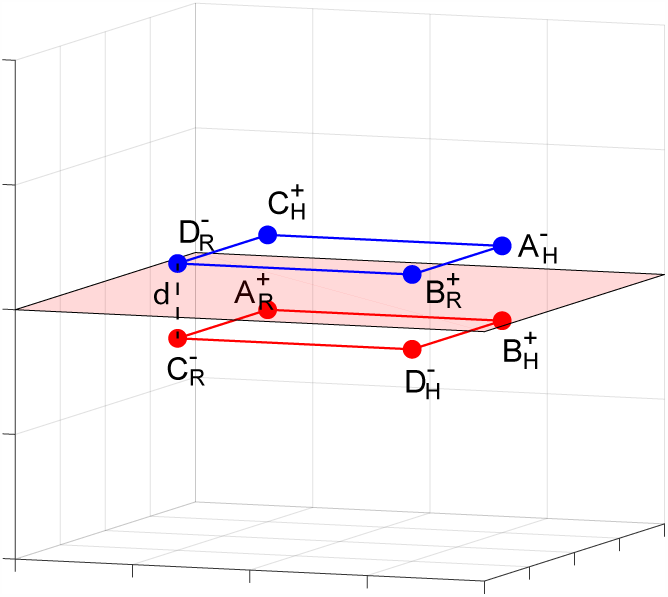
An example in which the clustering analysis fails to detect that context is in an abstract format. The firing rate space is represented as in Figure S6a. The points are arranged in a rectangular cuboid (a cube squeezed along the vertical direction), and the distance between the two squares that represent the two contexts is *d*. This distance is smaller than the sides of the two squares. When *d* is sufficiently small, the abstraction index is one or less than one, despite the fact that context is in an abstract format according to the CCGP (see the separating hyperplane which illustrates one linear classifier that clearly allows for cross-condition generalization). This type of geometry (the squeezed cube) may be observed whenever variables are encoded with different strengths (see also text in this section).

This should also clarify that, despite the partial homonymy, CCGP is not a synonym of generalization performance, the quantity that is the central object of statistical learning theory. In fact, generalization performance typically refers to testing the classifier on new data that is sampled from the SAME conditions used at training, while CCGP (it is important to stress) tests the classifier on new data from NEW conditions.

Analogously, quantities that are the purview of statistical learning theory and the empirical risk minimization approach in particular might seem related to our measure of abstraction to a deceptively high degree. For instance, the VC-dimension of a class of, say, maximum margin classifiers provides a PAC bound on their generalization performance as a function of sample complexity and training performance. So, it might seem superficially related to CCGP. What is important to keep in mind, however, is that we are not operating under the classical assumptions of statistical learning theory, as we don’t have the same goals. In particular, we’re are not interested in assuming that the held out response patterns are sampled from the same distribution as the training patterns (the so-called sampling hypothesis). On the contrary, we are assuming that the held out response patterns are being generated from unsampled experimental conditions, previously unseen by the classifier. And what we care about is measuring classification accuracy despite the covariate shift resulting from being presented with new experimental conditions. In other words, as profusely explained in the paper, we view abstraction as being supported by whatever geometrical property will guarantee generalization in previously unseen experimental conditions. Accordingly, if one really wanted to find a parallel between CCGP in our experiments and generalization performance within a statistical learning framework, it would be in the specific setting of domain adaptation, which investigates the classification performance in a target domain of classifiers trained in a different source domain. This is a theoretical connection that might be worth exploring in future work.

#### Does dimensionality of the neural response vectors trade off with our measures of abstraction?

Here we show that a high degree of abstraction does not necessarily entail low dimensionality. At first sight, it would however seem intuitive that the opposite is true. In fact, the class of neural geometries we denote as abstract require neurons to (at least approximately) exhibit linear mixed selectivity ^13, 14^, which would imply that these neural representations have low dimensionality. This is because the dimensionality of such a geometry would be equal to the number of the (in our case binary) abstract variables involved (not counting the possible offset from the origin of the neural activity space). This dimensionality is small compared to the total number of conditions (minus one), which corresponds to the maximal possible dimensionality achieved if all neural response vectors are linearly independent. In the idealized case in which all abstract variables have a parallelism score of one, the representations of the different conditions coincide with the vertices of a parallelotope, and it is easy to see that the dimensionality equals the number of variables, which is much lower than the maximal dimensionality.

It has been argued ^13, 14^ that high-dimensional neural representations are often important for flexible computation, because a downstream area may in principle have to read out an arbitrary dichotomy of the different conditions (i.e., an arbitrary binary function) to solve different tasks. This can be achieved using simple linear classifiers (used to model the type of computation that a readout neuron may be able to implement directly) only if the dimensionality of the neural representation is maximal. This desire for flexible computations afforded by high dimensionality seems to be in direct opposition to the low dimensionality implied by the abstract neural geometries that allow for cross-condition generalization.

However, there is in fact a large class of geometries that combine close to maximal dimensionality with excellent generalization properties for a number of abstract variables (which form a preferred subset of all dichotomies). Maximal dimensionality implies decodability by linear classifiers of almost all dichotomies. We will refer to this classifier-based measure of dimensionality as the ‘shattering dimensionality’. This quantity is similar to the one introduced in ^13^, where it was measured to be maximal in neural representations in monkey pre-frontal cortex. The geometries with high dimensionality and excellent generalization don’t have unit parallelism scores for the abstract variables (they are not exactly factorizable). A simple way to construct examples of such geometries is to start from a simple factorizable case, namely a cuboid, and then distort it to reduce the parallelism score.

We illustrate this cuboid geometry in the case of eight conditions. Here there are at most three completely abstract variables, as in our experimental data. We can generate an artificial data set with the desired properties by arranging the eight conditions initially at the corners of a cube (with coordinates plus/minus one), embedding this cube in a high-dimensional space by padding their coordinate vectors with zeros (here we use *N* = 100 dimensions, so we append 97 zeros), and acting on them with a random (100-dimensional) rotation to introduce linear mixed selectivity. We then distort the cube by moving the cluster center of each condition in a random direction - chosen independently and isotropically - by a fixed distance, which parameterizes the magnitude of the distortion. This operation reduces the parallelism score for the three initially perfectly abstract variables, which correspond to the three principal axes of the original cube, to values less than one. We sample an artificial data set of 1,000 data points per condition by assuming an isotropic Gaussian noise distribution around each cluster center.

On this data set we can run the same analyses as on our experimental data. In particular, we compute the parallelism score and the cross-condition generalization performance averaged over the three preferred (potentially abstract) dichotomies, with the results of these analyses shown in Fig. S8. In addition to these quantities, we compute the average decoding performance (ADP) across all 35 balanced dichotomies, which is a quantity related to the shattering dimensionality. Since the upper bound of ADP is one, we can think if it as a normalized measure of dimensionality. For small noise, the mean parallelism score of the three abstract variables starts very close to one and from there decreases monotonically as a function of the magnitude of the distortion. The same is true for their mean CCGP, but its decline is much more gradual and almost imperceptible for small distortions. This means that for intermediate values of the displacement magnitude (of order one), we still see excellent generalization properties across conditions, and the three preferred variables are still in an abstract format.

**Figure S8:**
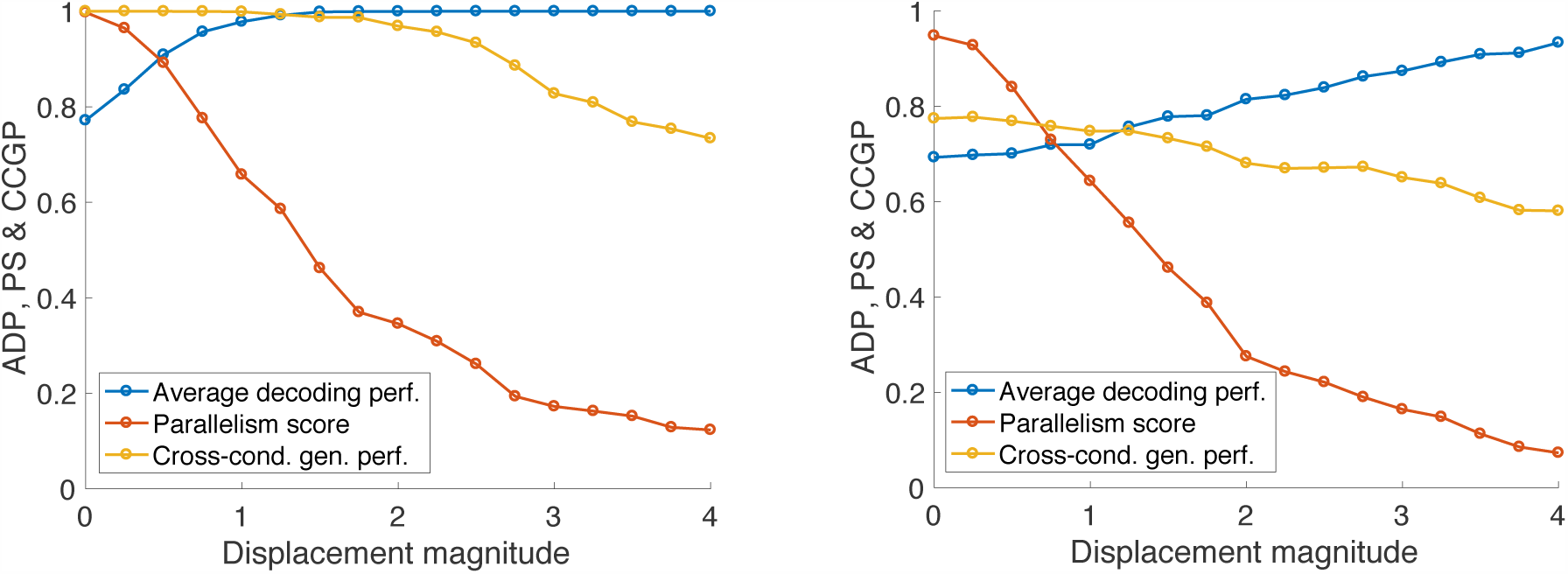
Analysis of an artificial data set generated by randomly embedding a (three-dimensional) cube in a 100-dimensional space, displacing its corners in independent random directions by a certain distance (displacement magnitude), and then sampling data points corresponding to the eight experimental conditions from isotropic Gaussian distributions around the cluster centers obtained from this distortion procedure. Average decoding performance (ADP, blue) is plotted across all 35 balanced dichotomies, which is a normalized measure of the dimensionality of the representation, as well as the mean PS (red) and CCGP (yellow) of the three potentially abstract variables for low (left) and high noise (right), with noise sampled as i.i.d. unit Gaussian vectors multiplied by overall coefficients 0.2 and 1.0, respectively. For low noise, both ADP and CCGP can simultaneously be close to one, indicating maximal dimensionality and the presence of three abstract variables for these representations. In the case of high noise, we observe a smooth tradeoff between CCGP (abstraction) and ADP (dimensionality).

In contrast, the ADP increases as a function of the magnitude of the distortion, due to the increasing dimensionality of the representation. Balanced dichotomies include the most difficult (least linearly separable) binary functions of a given set of conditions. If all these dichotomies can be decoded by linear classifiers, the dimensionality of the neural representation will be maximal. For small values of the displacement magnitude (for which the neural geometry is three-dimensional), some dichotomies are clearly not linearly separable. Therefore, ADP is initially substantially less than one, but it steadily increases with the degree of distortion of the cube. Crucially, it reaches its plateau close to the maximal value of one before the CCGP drops substantially below one. This indicates that there is a parameter regime in which this type of neural geometry exhibits almost maximal dimensionality, which enables flexible computation, but at the same time the geometry also enables abstraction (in the form of excellent CCGP) for the three preferred variables. Therefore, we can conclude that these two favorable properties (high dimensionality and high CCGP) are not mutually exclusive.

In the case of larger noise, the PS and CCGP start out at values substantially smaller than one already for zero distortion. Although the qualitative trends of all the quantities discussed remain the same, the tradeoff between ADP and CCGP (i.e., between flexible computation and abstraction) is much more gradual under these circumstances.

### S3 Selectivity and abstraction in a general linear model of neuronal responses

We model the selectivity of a neuron to the various task variables in a given time bin as FR with the following linear regression model (see also Supplementary Material of ^13^):

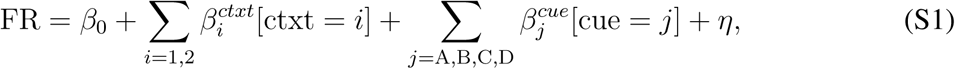

where the terms in square brackets represent a design matrix whose elements indicize the type of trial according to context type and cue identity. The coefficient *β*_0_ is a bias and *η* represents a Gaussian noise term. Fitting the model described by equation (S1) to the firing rate of a neuron is related to performing a 2-way ANOVA, where the two factors correspond to context type, and cue identity. The *β* coefficients that are determined by the fit quantify the selectivity to their corresponding aspects. Since equation (S1) does not include any interaction term among the factors, the model is linear. Because of this, the corresponding neuron would be denoted as a *pure selectivity neuron* neuron if only *β*’s corresponding to one factor are significantly non-zero, or it would be denoted as a *linear mixed selectivity neuron* in case at least one *β* for each factor is significantly non-zero (see ^13^).

The model (S1) can be complemented with additional factors aimed at fitting the deviation of the activity FR from a linear model. These are typically included as 1-factor interaction terms:

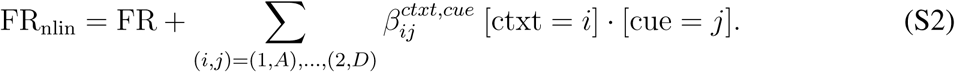

Whenever any coefficient of an interaction term is significantly different from zero, the neuron is denoted as a *nonlinear mixed selectivity neuron*. Nonlinear interaction terms increase the dimensionality of the neural response patterns, which benefit the downstream linear decoding^13^. However, increasing dimensionality is generally at odds with abstraction as discussed in Section S2.

Linear mixed selectivity on the other hand does not increase dimensionality beyond the dimensionality of a population of pure selectivity neurons that collectively display all the factors explaining the activity of the pure selectivity neurons (see ^13^).

### S4 The recorded neurons are not highly specialized

An ensemble of neurons could in principle encode multiple abstract variables simply by assigning each neuron to be tuned to precisely one of these variables. In this case, the ensemble can be divided into a number of subpopulations each of which exhibits pure selectivity for one of the abstract variables. Geometrically, this situation is similar to the one depicted in Fig. 3a. The situation in which neurons exhibit mixed selectivity can be obtained by rotating this geometry, as shown in Fig. 3b. This rotated representation would show the same generalization properties. In the data, we do not observe many pure selectivity neurons. If neurons were highly specialized for particular variables, training a linear classifier to decode that variable should lead to very large readout weights from the associated specialized subpopulation, but very small readout weights from other neurons (which might specialize on encoding other variables). Therefore, in a scatter plot of the (absolute values) of the readout weights for different classifiers we would expect specialized neurons to fall close to the axes (with large weights for the preferred variable, but small ones for any others). If this situation occurred for a large number of neurons, we might expect a negative correlation between the absolute values of the decoding weights for different variables. However, in fact the weights do not cluster close to the axes, as shown in Fig. S9. The correlation coefficients of their absolute values are positive. We conclude that highly specialized neurons are not particularly common in the neural ensembles we recorded, i.e. pure selectivity appears to be no more likely than coding properties corresponding to a random linear combination of the task-relevant variables.

For each neuron we can consider the three-dimensional space of the readout weights for the three variables (components of three different unit-norm weight vectors in the space of neural activities). Clearly neurons with large readout weights for the linear classifier trained to read out a particular variable are important for decoding that variable. Some neurons have larger total read-out weights than others (e.g. we can consider the sum of squares of the three weight components, corresponding to the squared radius in the three-dimensional space) and are therefore more useful for decoding overall than others which have only small readout weights.

We can also rank neurons according to the angle of their readout weight vector from the axis associated with one of the variables in the three-dimensional space. This quantifies the degree of specialization of the neuron for the chosen variable. We call the absolute value of the cosine of this angle the pure selectivity index. We can now ask whether neurons with a large pure selectivity index are particularly important for generalization, as quantified by the cross-condition generalization performance (CCGP). This can be tested by successively removing neurons with the largest pure selectivity indices from the data and performing the CCGP analysis on the remaining population of (increasingly) mixed selectivity neurons. The results of this ablation analysis are shown in Fig. S10, in which we plot the decay of the CCGP with the number of neurons removed. It demonstrates that while pure selectivity neurons are important for generalization (as expected in a pseudo-simultaneous population of neurons with re-sampled trial-to-trial variability, in which the principal axes of the noise clouds are aligned with the neural axes), they are not more important than neurons with overall large decoding weights which typically have mixed selectivity.

We can perform the same analyses also for the neural network model of Fig. 5. We focus on the neural representations obtained from the second hidden layer of the trained network. Fig. S11 shows a scatter plot of the decoding weights for the parity and magnitude variables. As in the neural recordings, many neurons exhibit mixed selectivity. We can again successively ablate neurons with the largest pure selectivity indices. Unlike for the case of the neural data, this only has a small effect on CCGP (see Fig. S11). Presumably this is because here the neural activities are simultaneously ‘recorded’, and therefore the decoder can use information contained in the correlations between different neurons in these representations. Unlike the case of resampled noise in a pseudo-simultaneous population, the principal axes of the noise clouds are not necessarily aligned with the neural axes. For such a simultaneously observed representation there is no reason why pure selectivity neurons should contribute preferentially to the generalization ability of the network.

**Figure S9:**
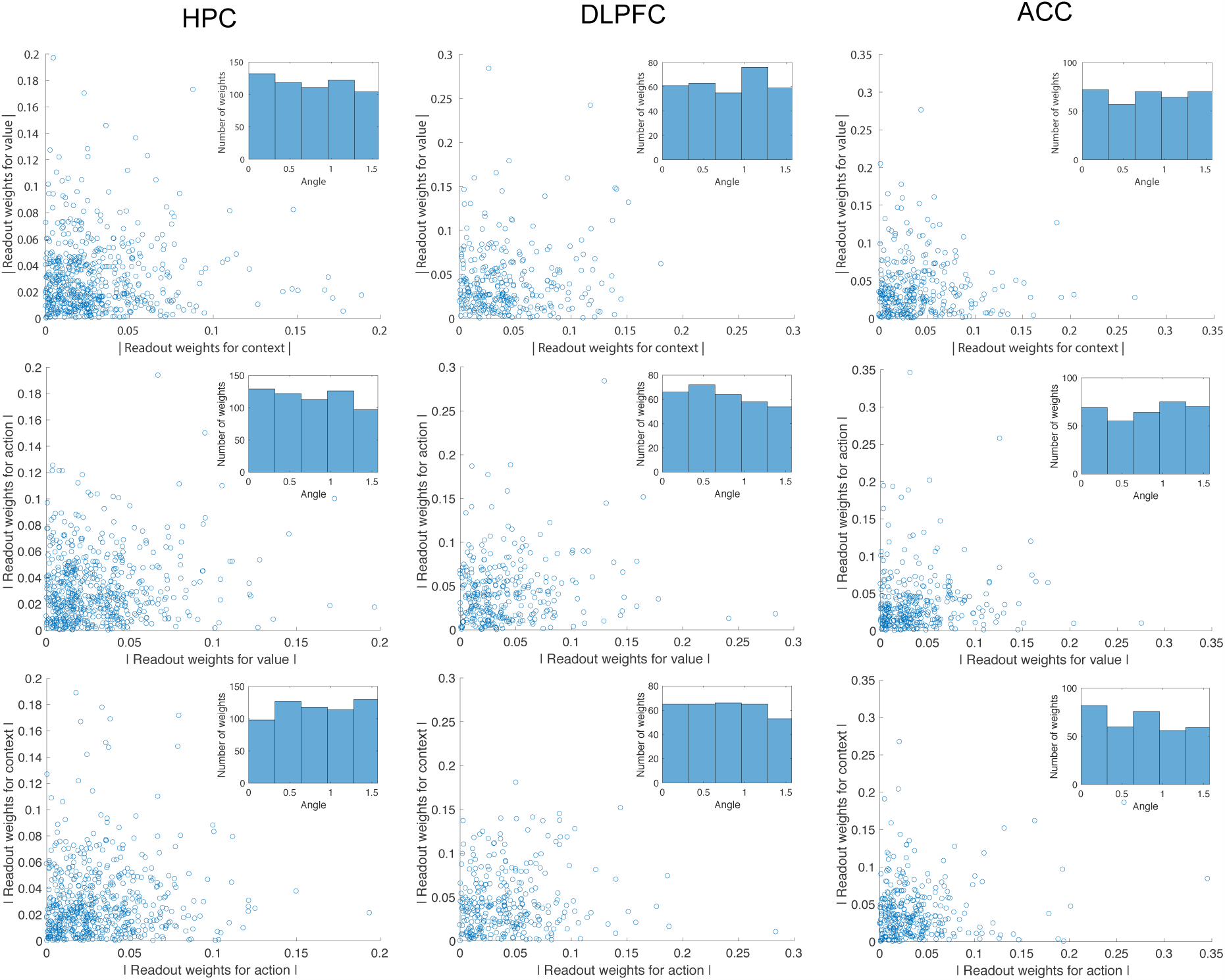
Two-dimensional scatter plots of the absolute values of the (normalized) decoding weights for the three task-relevant variables. The three columns (from left to right) correspond to HPC, DLPFC, and ACC. The three rows show the magnitudes of the weights for pairs of variables plotted against each other: context vs. value (top), value vs. action (middle), and action vs. context (bottom). The inset in each scatter plot shows a histogram of the weight counts as a function of the angle from the vertical axis (in radians). These distributions are approximately uniform, and therefore pure selectivity neurons (whose weights would fall close to one of the axes in the scatter plots) are not prevalent. Similar distributions have been observed in the rodent hippocampus 47.

**Figure S10:**
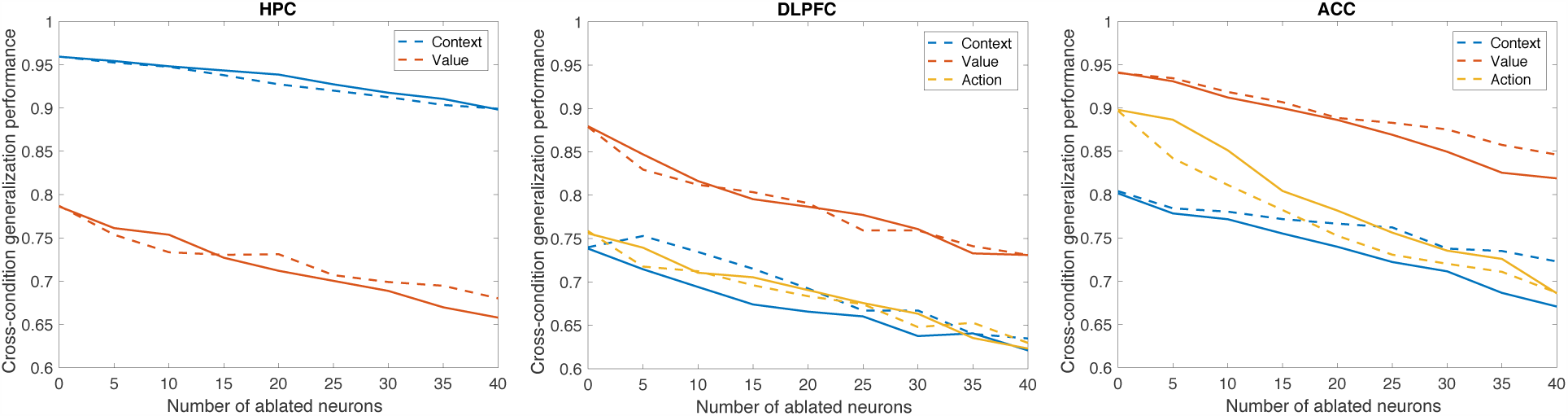
CCGP as a function of the number of ablated neurons for the HPC (left), DLPFC (middle), and ACC (right). The solid lines show the decay of CCGP if we successively remove the neurons with the largest pure selectivity indices for context (blue), value (red) or action (yellow). The dashed lines show the decline of the CCGP for the same three variables if we instead ablate neurons with the largest sum of squares of their three decoding weights (i.e., those with the radial position furthest from the origin in their three-dimensional weight space), independent of their pure selectivity indices. The two sets of curves are rather close to each other, and thus these two sets of ablated neurons are of similar importance for CCGP. (For HPC, the CCGP of the action variable is always below chance level for both curves; not shown). This is similar to what has been observed in simulations of deep networks 48.

### S5 Dimensionality of the neural representations

We utilize a technique developed in ^49^ to estimate a lower bound for the dimensionality of the neural response vectors in a specific time bin during a task. Similarly to our other analyses, we build average firing rate patterns for all recorded neurons by averaging spike counts sorted according to task conditions indexing the trial where the activity is recorded (current trial) or the previous trial. The spike counts are z-scored and averaged in 500 ms time bins displaced by 50 ms throughout the trial. We then apply the method presented in ^49^ on the obtained average firing rate activity patterns independently within each 500 ms time bin. This procedure allows us to bound the number of linear components of the average firing rate patterns that are due to finite sampling of the noise, therefore providing an estimate of their dimensionality.

Figure S12 shows the result of this analysis for all neurons recorded in HPC, DLPFC and ACC for which we had at least 15 trials per condition. For average firing rate patterns obtained by sorting spike counts according to the 8 conditions of the current trial (continuous lines), dimensionality peaks at its maximum possible value shortly after the presentation of the image for all three areas. The dimensionality for firing rate patterns obtained by sorting the activity according to the condition of the previous trial remains around 5 throughout the trial, which is close to the value to which dimensionality in the current trial decays towards the end of the trial.

**Figure S11:**
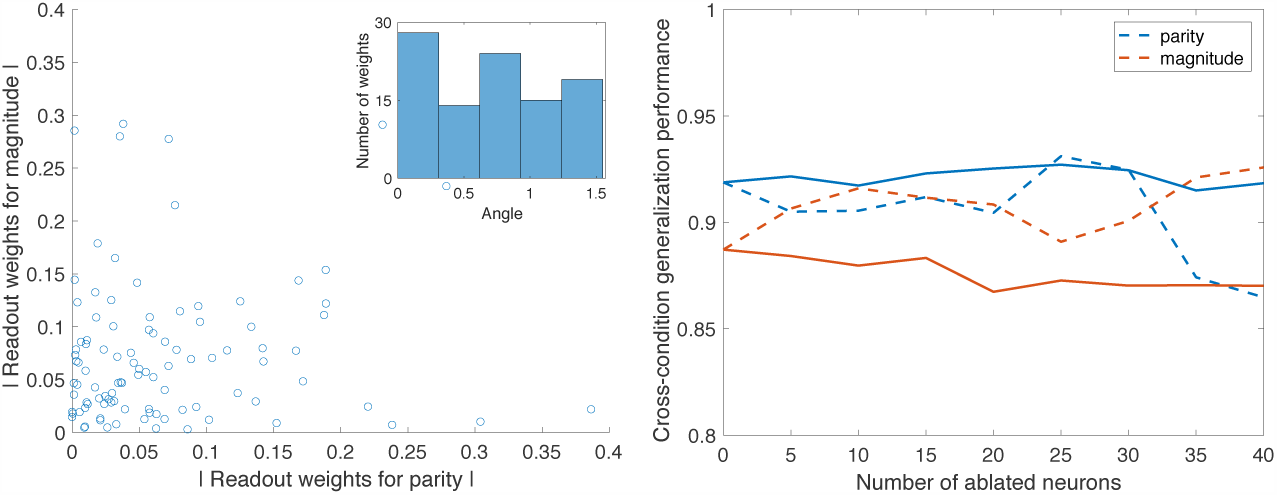
Specialization and ablation analyses of second hidden layer in the neural network of Fig. 5. Left: Two-dimensional scatter plot of the absolute values of the (normalized) decoding weights for the parity and magnitude dichotomies, as in Fig. S9. Right: CCGP when training on three digits from either side of the magnitude (red) and parity (blue) dichotomies and testing on the fourth one as a function of the number of ablated neurons. We ablate the neurons with the largest pure selectivity indices (solid lines), or the ones with the largest sum of squared decoding weights (dashed lines), as in Fig. S10. Note that even though we dont z-score the neural representation for the computations of the CCGP, or for any of the analyses shown in Fig. 5, the decoding weights shown here (and used to select the ablated neurons) are taken from classifiers trained on z-scored representation, since otherwise they would hardly be indicative of the relative importance of different neurons for decoding.

**Figure S12:**
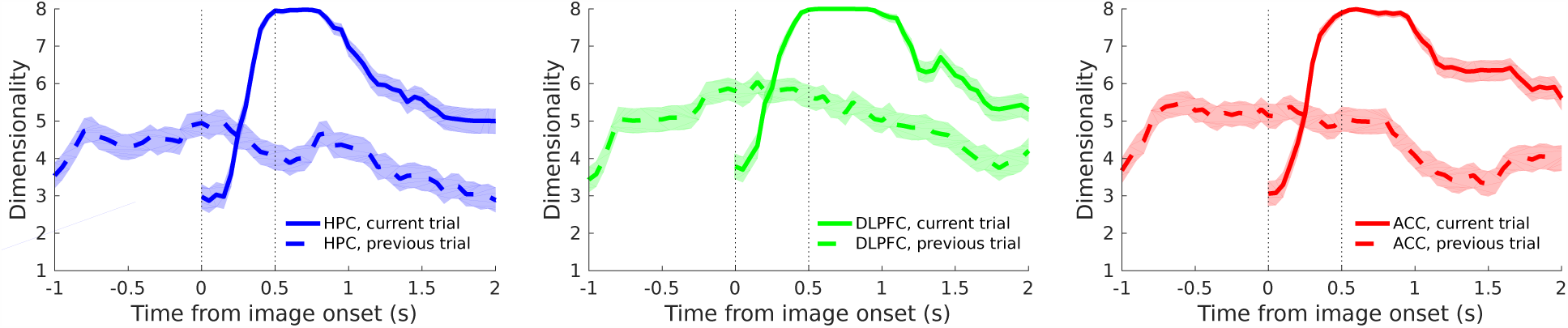
Dimensionality of the average firing rate activity patterns as a function of time throughout the trial. The left panel illustrates the result of the analysis developed in 49 on HPC, the central panel refers to DLPFC, and the right panel to ACC. Continuous lines refer to the analysis carried out on average firing rate patterns obtained by averaging spike counts according to the task conditions of the trial that was being recorded (current trial). The dashed line shows the same but for conditions defined by the previous trial. The lines indicate the number of principal firing rate components that are larger than all noise components, averaged over 1000 re-samplings of the noise covariance matrix (see 49). The shadings indicate the 95% confidence intervals estimated using the method for quantifying uncertainty around the mean of bounded random variables presented in 50.

### S6 Deep neural network models of task performance

The study of performance-optimized neural networks is becoming increasingly common practice in computational neuroscience. This technique consists in training an artificial neural network model on the simulated version of a behavioral task of interest, and examining the neural activations generated by the neural network during or after learning the task. Following such practice, we train neural networks to perform a simulated version of our task under two learning paradigms and we show that after learning the neural representations are similar to those recorded in the experiment. In particular, we show that the learning algorithms that we use both lead to neural representations in which the hidden variable context is in an abstract format, and multiple abstract task-relevant variables are simultaneously represented. We assume that the inputs provided to the neural network are the following observable quantities: the stimulus, action and outcome in the previous trial, and the stimulus in the current trial. It is therefore intriguing that through the process of learning to execute the context-dependent task, the network learns to internally represent context (a latent variable not explicitly represented in the task), and moreover that this representation is in an abstract format as quantified by our metrics. The functional role of this abstract context representation is to modulate the internal representation of the current stimulus, as demonstrated by the fact that the trained network selects the correct action in response to a stimulus in each given context.

As a first general learning paradigm we investigate *Reinforcement Learning*: our neural network model simulates the behavior of the animals engaged in the task and is trained by trial and error to maximize its performance on the simulated task. We then take a drastic simplification step and model the interaction between the animal and its environment during the task as a *supervised learning* paradigm: our network model is essentially trained to output the correct (assumed to be known) behavioral response and predict the resulting outcome, given the current stimulus, and the sequence of stimulus, correct response and outcome in the previous trial. As we will see, this simplification still preserves the main features of the activity patterns generated by the more complex Reinforcement Learning model, while at the same time providing us with direct control of the statistics of the inputs presented at training (which are no longer dictated by the behavior of the model itself), thereby enabling us to systematically study their effect on the abstraction of the variables emerging during learning of the task.

#### Reinforcement Learning model

We simulate a sequential version of the task that is used as an environment to train by Reinforcement Learning a decision policy (an agent) parametrized as a multi-layer network with two hidden layers (whose size is indicated in Figure S13a), three outputs (one for each of the actions available to the agent), and ReLU activations.

In order to simplify the modeling, we decided to explicitly encode as external inputs all the information necessary to determine the current context. In particular, in addition to the current stimulus, we also provide to the network the previous stimulus, the previous action, and the resulting previous outcome. Previous stimulus, action and outcome are necessary and sufficient to determine the current context (assuming there was no switch between the previous and current trial). One might postulate that these inputs are internally maintained through, for instance, some recurrent synaptic circuit. However, representing the inputs explicitly does not qualitatively affect the model, besides allowing us to eschew a recurrent architecture in favor of a simpler feed-forward one. Notice, moreover, that even if the model were to directly maintain an encoding of context, information about the outcome in the previous trial (together with previous stimulus and action) is essential in order to correctly predict the correct action after a contextual switch. As we will detail in the next paragraphs, the result of this modeling exercise is that, indeed, after training, our model internally encodes the current context when previous stimulus, action and outcome are supplied to the network. Moreover, context is being represented in an “abstract format” (high CCGP and parallelism score), and this modulates the hidden layer representation of the current stimulus to result in the selection of the correct context-dependent action.

**Figure S13:**
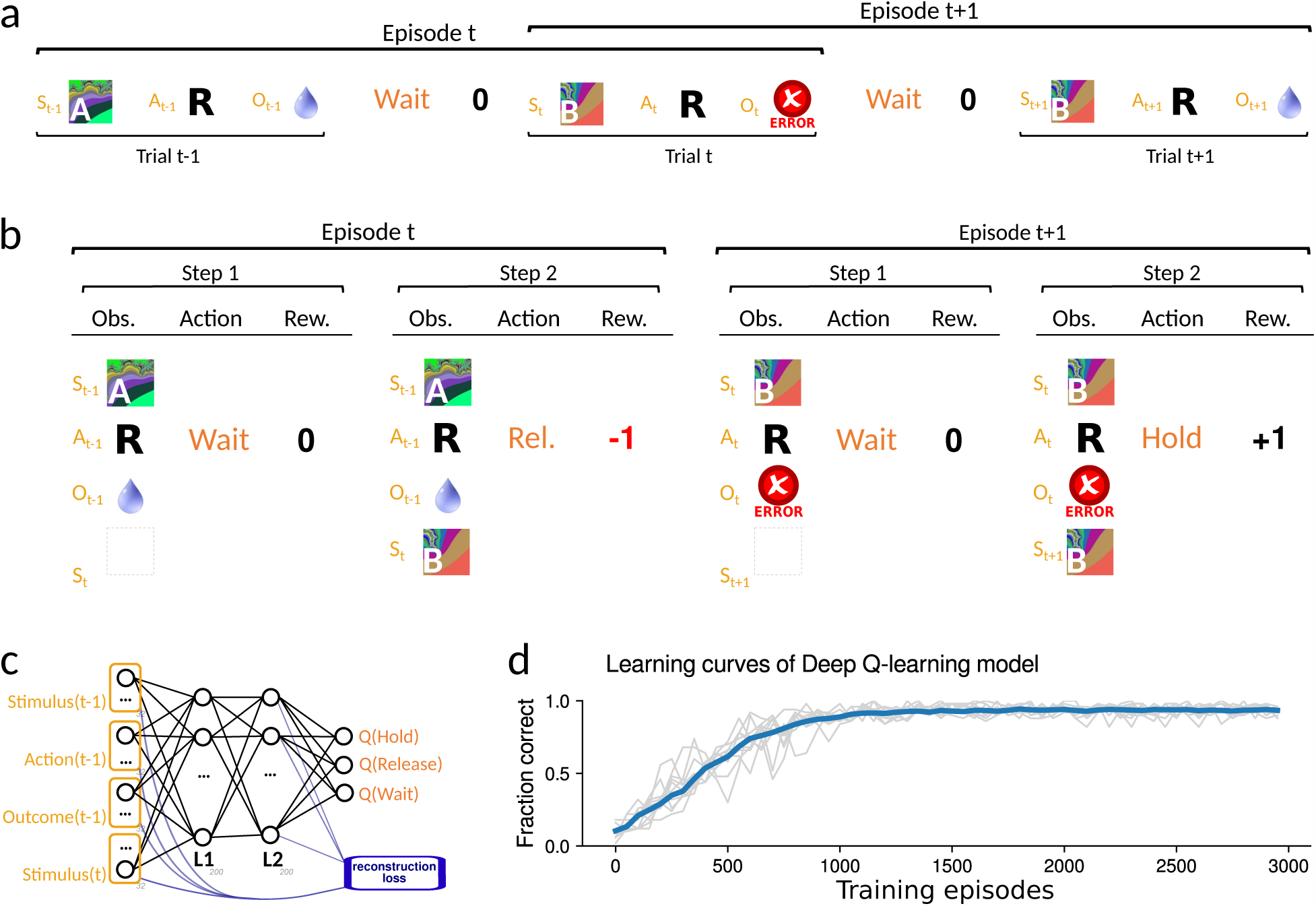
Reinforcement Learning (RL) model trained with Deep Q-learning (DQN) on a simulated version of the serial reversal-learning task. a., b. An episode in the simulated environment is composed of two sequential steps, each corresponding to one trial. In the first step, corresponding to the pre-stimulus interval in the trial, the network receives as observations the stimulus, the action and the outcome in the previous trial, at which point it is required to issue the action “Wait” to initiate a new trial. In the second step of the episode the network receives the current stimulus (in addition to all observations presented in the first step) and is required to issue the correct action in the task (“Hold” or “Release”). If at any step the agent issues an incorrect action, it receives a reward of −1 and the episode terminates. Otherwise, it receives a reward of 0 after correctly issuing “Wait” at the end of the first step, and for trials in which the agent issues the correct action, it receive a reward corresponding to the value of the stimulus in the task at the end of the second step (+1 or 0 depending upon whether it is a rewarded trial type). c. The agent is parametrized as a two-hidden layers neural network trained with DQN. We add a reconstruction loss to the optimization loss (indicated in blue in the lower right corner of the panel) that forces the network to find representations in the L2 hidden layer that can linearly reconstruct the inputs (i.e., the reconstruction loss is the mean squared error between the inputs and a linear transformation of the L2 activations). d. Learning curves for the DQN model on the serial-reversal task. The blue line is the fraction of correct actions averaged over 100 random initializations of the network and blocks of 50 episodes. The gray lines show the performance of 10 randomly chosen individual networks (also in blocks of 50 episodes).

In constructing the model, all observable inputs (information observable in the environment) are encoded and presented to the network as binary input vectors with added Gaussian noise sampled independently for every trial. Figure S13a details the construction of these input vectors. At the beginning of a simulation we randomly sample 32-dimensional binary vectors encoding each of the values of all features appearing in the task (previous stimulus, previous action, previous outcome and current stimulus). An input representing a given task condition is then composed by stacking the binary vectors corresponding to the features describing the condition, and adding centered Gaussian noise with amplitude of 0.4.

An episode in the environment consists of two steps. In a first step corresponding to the prestimulus interval in the task, the agent observes the stimulus, action and outcome of the previous trial. The correct action at this step is “Wait”, upon which the agent receives a reward of 0. If the agent mistakenly selects “Hold” or “Release”, it receives a negative reward of −1 and the episode is terminated. In the second step of each episode, in addition to the observables presented in the first step, the agent observes the stimulus of the current trial (Figure S13a). At this point the agent has to select the correct action between “Hold” and “Release”, which depends on the currently active context and the current stimulus identity. Upon selection of the correct action, the agent receives a reward of +1 for a rewarded trial, or a reward of 0 in the case of an unrewarded trial. Again, if the agent selects the wrong action, it receives a negative reward of −1. At this point the episode terminates, and a new one is initiated whose context is chosen to be identical to the one of the previous trial with a probability of 98%. Notice that information about the context identity is never made apparent in the observations, but is only reflected in the determination of the correct action, given the current stimulus. As a consequence, assuming that the context hasn’t changed between the previous and current episode, the current context can be inferred from the triplet of previous stimulus, previous action and previous outcome, since these uniquely determine the correct action in the previous episode and therefore its context.

The particular choice of dividing each episode into two steps overlapping two adjacent trials was dictated by the requirement of being able to analyze a trial epoch corresponding to the pre-stimulus interval analyzed in the recorded data. Moreover, explicitly providing information about the previous trial in the input to the network allows us to eschew complicated mechanisms to preserve information across time steps, such as for example having to resort to a recurrent neural network model. Future studies will consider these more complicated mechanisms. The environment modeling choices just described thereby allow us to model the agent with a very simple feed-forward architecture, as depicted in Figure S13a. The only slightly unconventional element in our policy network architecture is the addition of a *reconstruction loss* (Figure S13a, lower right). This consists in a weighted term added to the optimization cost that is proportional to the squared difference between the input to the networks and a linear transformation of the activations of the second hidden layer. The role of this additional cost is to allow us to tune the amount of information that the hidden layer representations encode about the inputs. Practically speaking, this provides a mechanism to counteract a tendency observed in our simulations: larger numbers of training epochs and increasing depth tend to be associated with the representations of previous outcome and action variables in a less abstract format (as measured by CCGP and PS). This effect is analyzed more quantitatively using the supervised learning model in the next section.

The policy network described in the previous paragraph is trained to select the correct action with Deep Q-learning^15^. In particular, we use a frozen copy of the policy network that is being updated only every 10 episodes to estimate the Q-values corresponding to an observation-actions-reward triplet randomly sampled from a replay memory buffer of 5000 steps in the environment. After every step, a plastic copy of the policy network is being trained to fit the Q-values estimated from the frozen policy network by using one step of adam-SGD^51^ (with the default parameters in PyTorch ^52^, aside from a learning rate of 0.005 with a decay factor of 0.999) and mini-batch of size 100. The use of a slowly updated frozen copy of the policy network to estimate the Q-values is a technique that had been proposed in ^15^, and it helps to stabilize the learning procedure. Moreover, as customary to promote exploration in Reinforcement Learning, actions are selected using an *ϵ*-greedy algorithm, i.e., they are selected based on the largest Q-value with a probability 1 −*ϵ*, and selected uniformly at random with probability *ϵ*. We set *ϵ* = 0.9 at the beginning of learning and decrease it exponentially to 0.025 with a time constant of 500 steps. Finally, we use a discount factor of *γ* = 0.99.

Figure S13d shows the average learning curve for a population of 100 policy networks initialized at random and trained on random instantiations of the environment described in the previous paragraphs. The blue curve shows the fraction of correct terminal actions within blocks of 50 episodes, averaged across 100 networks, as a function of episode.

The results of analyzing the activity of the Reinforcement Learning model are presented in Figure S14. All analyses are conducted during the first step in the simulated tasks, which emulates the pre-stimulus interval in the experimental task, on a population of 100 randomly initialized and independently trained networks. The figure shows the results of the analyses averaged over the population of models, both before (grey data points) and after training (orange data points). We compute decoding accuracy (first row of panels), PS (second row) and CCGP (third row) for context (first column), value (second column) and action (third column).

The first column of Figure S14 shows that the *context* cannot be decoded with above-chance accuracy (50%) in the input but can already be decoded with high accuracy starting from the randomly initialized first hidden layer (gray datapoints). This is consistent with previous results demonstrating that large enough networks of randomly connected neurons with non-linear activations generate linearly separable patterns with high-probability (see ^23, 53, 54^).

**Figure S14:**
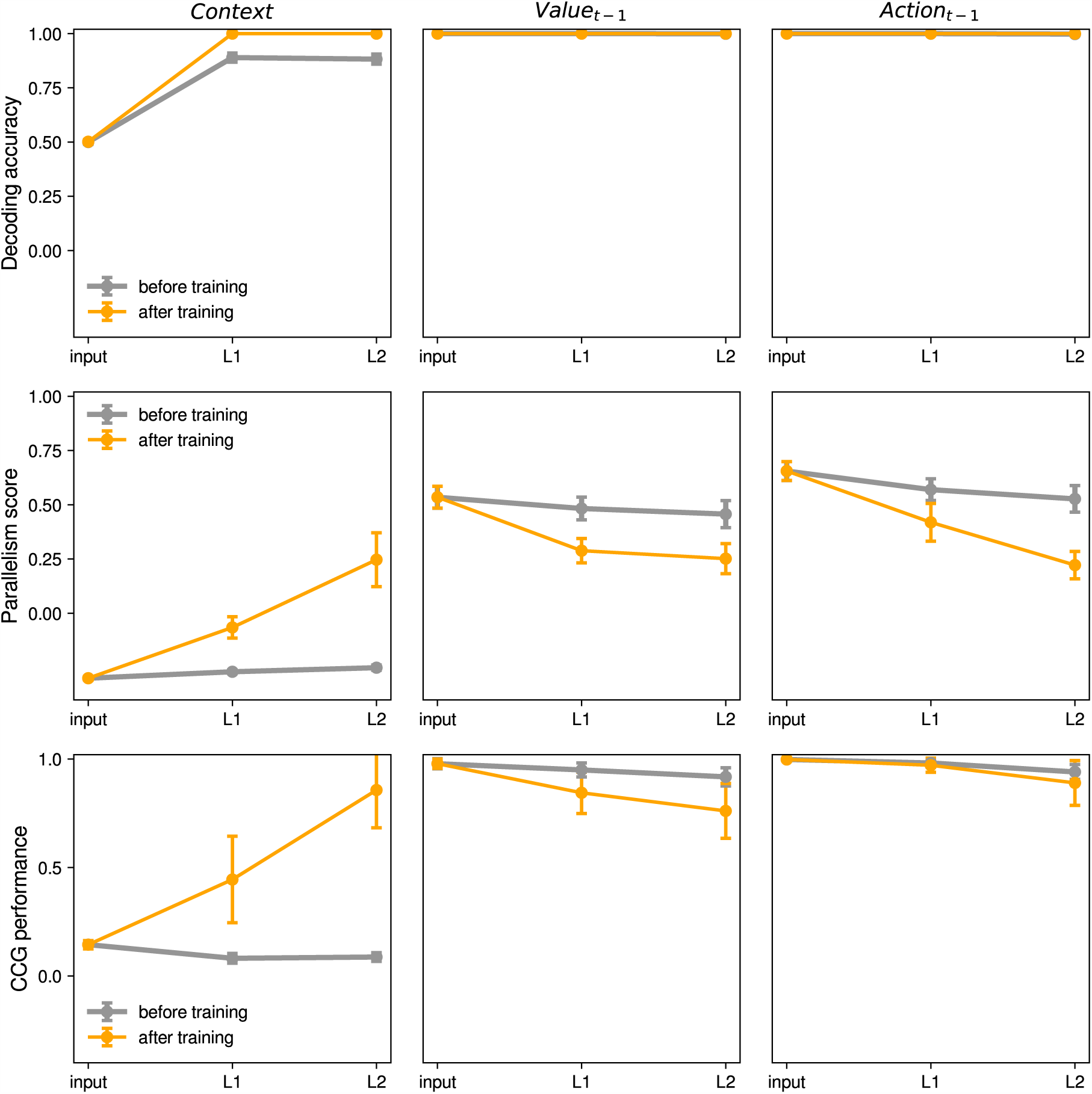
Analysis of the activity generated by the Reinforcement Learning model: *decoding accuracy* (first row), *parallelism score* (second row) and *cross-condition generalization performance* (third row) during the simulated first step (pre-stimulus epoch) in the task, for the variables *context* (first column), *previous value* (second column) and *previous action* (third column). In each panel we plot one of the quantities of interest as a function of the layer where it is measured in the architecture: inputs, first hidden layer (L1) and second hidden layer (L2) (see Fig. S13). The plots show the mean quantity of interest across 100 randomly initialized and independently trained networks, before (grey data points) and after training (orange data points). Error bars indicate standard deviations computed over the same distribution of 100 networks.

The second row of Figure S14 shows that, despite being decodable with high accuracy in both hidden layers, context only becomes appreciably abstract in layer L2, where on average across the models CCGP becomes significantly higher than chance (i.e., 50%). Correspondingly, the average PS in L2 is fairly high at around 0.25 (Figure S14, first plot in the second row). Moreover, in both hidden layers of the trained models, PS and CCGP for context are significantly higher than in untrained models. In this respect, it is worth emphasizing that for the variable context, the CCGP observed in randomly initialized models is consistently and considerably below chance level. This reflects the anti-correlation structure induced by the contextual character of the task: stimuli have to be associated to opposite context labels in the two contexts, which will naturally induce a linear decoder trained on a subset of conditions to predict the wrong context for the untrained conditions. This structure is also reflected in the parallelism score which tends to be negative for untrained models.

As for the previous value and action variables, they maintain relatively high CCGP and PS after training, but there was a general tendency for representations to become progressively less abstract as a function of training and depth of the layer. This presumably reflects the fact that value and chosen action in the previous trial are not features that are directly predictive of the current value and action, although they can be utilized to compute the current context and therefore provide information that could allow an agent infer the current value and action.

Besides comparing the PS and CCGP of the trained models with those of the untrained models as baseline, we can quantify their statistical significance using the more stringent statistical tests developed to analyze the experimental data. In particular, we can use the geometric random model as a null-distribution to establish the statistical significance of CCGP and use a shuffle test as a null-distribution for the PS (as in Figure 4). Accordingly, Figure S15 shows beeswarm plots of the probability that the CCGP and PS of each of the trained Reinforcement Learning models is above the distributions given by the geometric random model and a shuffle test, respectively, for all variables in the task individually (context, previous value, and previous action) and all variables simultaneously for the same model (All). These probabilities are empirically estimated over 100 trained models and 1000 samples of the null-distributions per trained model. Figure S15 also displays the mean of the null-distributions (dotted lines) and the 95th percentile (dashed line). Above each plot, the fraction of trained models whose PS and CCGP is above the 95th percentile of the null-distribution is depicted. Therefore, particularly in layer L2, a considerable fraction of trained models display a PS and CCGP above the 95th percentile of the null-distributions. For instance, in L2 40% of the trained models have a PS that is higher than 95% of the shuffle null-distribution simultaneously for all variables, while 11% have a CCGP above 95% of the null-distribution obtained with the random model.

#### Supervised Learning neural network model

Besides investigating a Reinforcement Learning model of the task, we also devised simulations in a much simpler open-loop learning paradigm:

**Figure S15:**
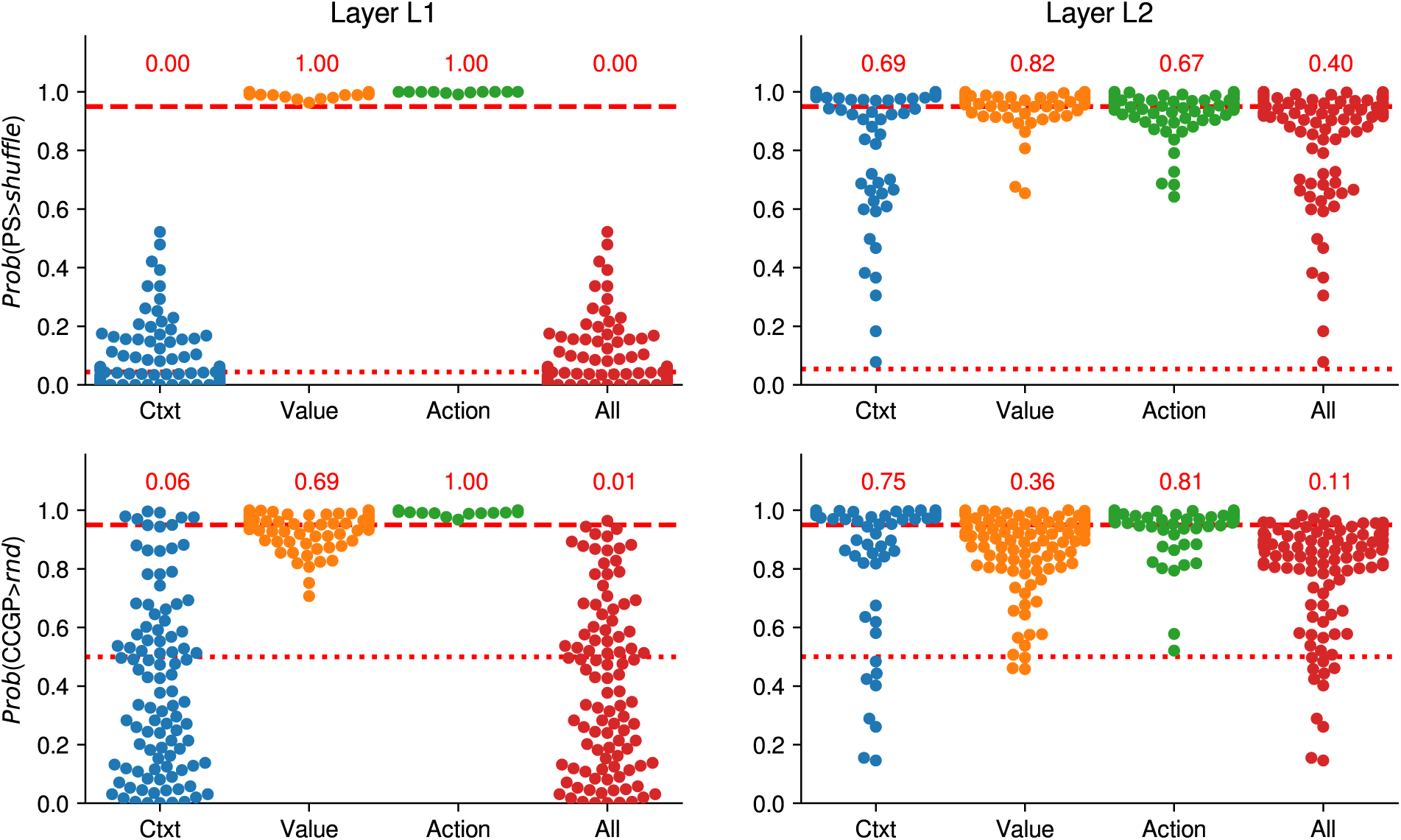
Beeswarm plots of the probability of PS and CCGP of trained Q-learning models to be above the null-distributions given by the geometric random model and a shuffle test of the data (empirically estimated with 1000 samples per model), respectively (see Figure 4). Every dot in the graph represents a trained model. The first row of plots shows the probability that the Parallelism Score of each model for every variable individually (Context, Value and Action) and all variables simultaneously (All) is above the null-distribution given by a shuffle of the data in layer L1 (first column) and L2 (second column). The second row is laid out similarly as the first one, but shows the probability that the CCGP of each model is above the null-distribution given by the geometric random model. The dotted lines correspond to the mean of the null-distribution, and the dashed lines correspond to the 95th percentile of the null-distribution. The numbers written in red above each plot report the fraction of models that are above the 95th percentile threshold.

**Figure S16:**
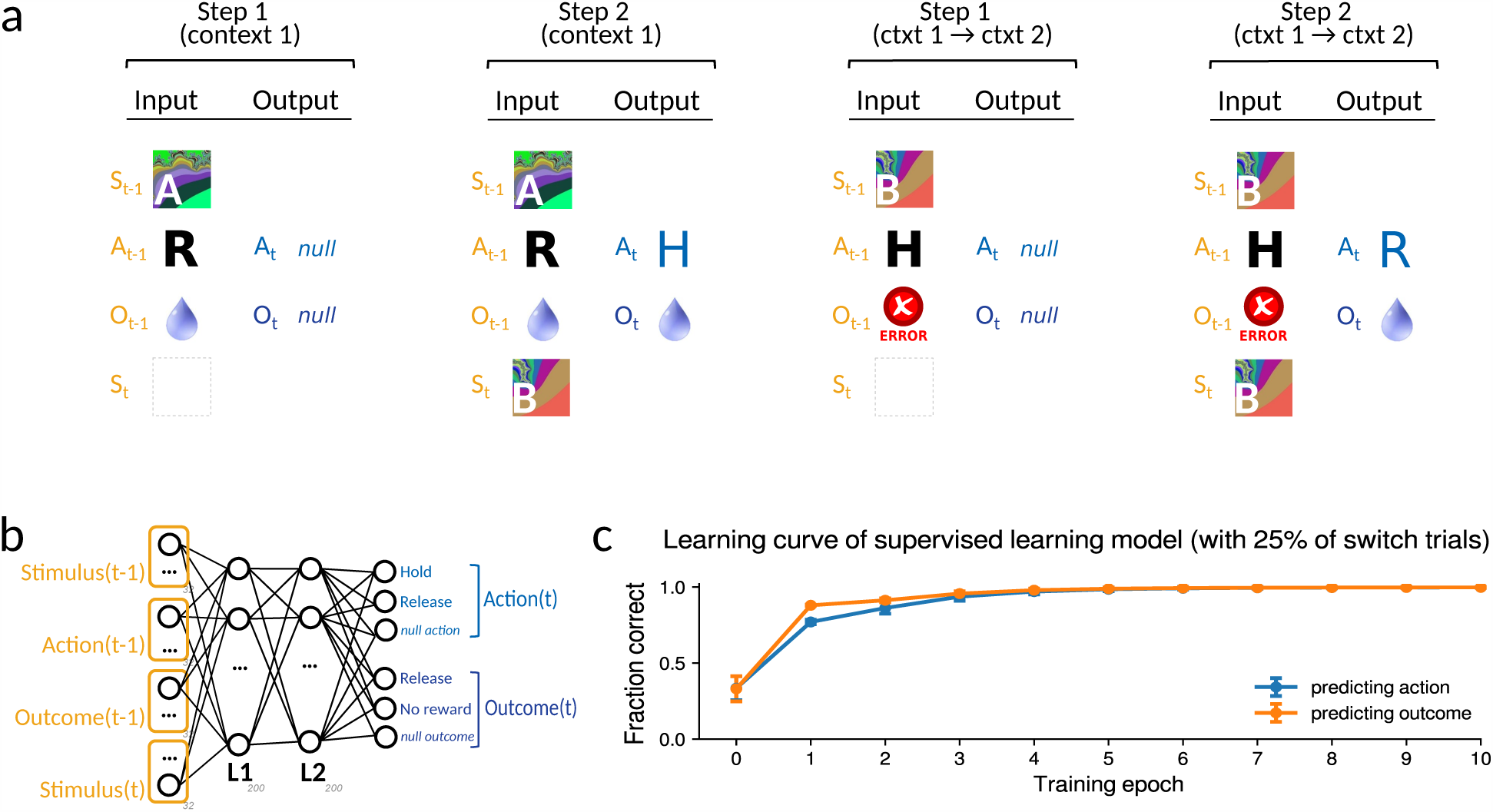
Supervised learning model. a. The supervised learning model is trained on two types of input-output combinations, corresponding to sequential steps in a trial. The inputs for the first step (the pre-stimulus interval in the trial) are composed of the stimulus, the action and the outcome (reward) in the previous trial. The target output for the first step is a ‘null’ action and outcome value (corresponding to a third null action and a third null outcome neuron). Inputs corresponding to the second step encode the current stimulus (along with the features that were already presented in the first step), and the second step output encodes the correct action (Hold or Release) and outcome (Reward or No reward) in the trial. Besides being distinguished by whether they correspond to the first step or second step in the trial, generated inputs can be distinguished depending on whether or not they correspond to a switch trial. Inputs generated from non-switch trials (see first and second input-output combinations in the panel) define the correct action and subsequent outcome, under the assumption that the current context is the same as in the previous trial (encoded in the input). Inputs generated from switch trials (the first trial after a context switch) are characterized by outcome features encoding an error in the previous trial (see second and third input-output combinations in the panel), implying that the observed stimulus-action combination is incorrect and counterfactually defining the currently correct context and action. b. The neural network is a multi-layer network with two hidden layers (number of unites indicated in the figure) trained with backpropagation in a supervised way to output the correct action and outcome. c. Learning curves for the (simultaneously learned) action and value prediction tasks at the end of the second step (corresponding to the response and value prediction of the animal) as a function of training episodes. Every epoch consists of 128 mini-batches of 100 noisy versions of the inputs just described. Data points are the means and standard deviations across 100 distinct random initializations of the network.

supervised learning. In this case, the network is trained to associate inputs corresponding to the stimuli presented to the monkeys with outputs corresponding to the correct action and expected outcome. Given a task condition, the inputs are generated in the same way as for the Reinforcement Learning model. At the beginning of a simulation we randomly sample 32-dimensional binary vectors encoding each of the values of all features appearing in the task (previous stimulus, previous action, previous outcome, current stimulus, correct action and predicted outcome). An input representing a given task condition is composed by stacking the binary vectors corresponding to the features describing the condition, and adding centered Gaussian noise with amplitude of 0.4. A corresponding output is built by stacking the binary vectors encoding the correct action and predicted outcome in the current condition (see Figure S16). As for the Reinforcement Learning case, we simulate two task epochs for each trial type: a pre-stimulus epoch defined by the previous outcome, previous action and previous stimulus (the activity of the input neurons encoding the current stimulus is set to zero), and a second epoch where also the current stimulus is presented. For inputs corresponding to a pre-stimulus epoch the network is trained to output a ‘null’ action by activating a third action neuron, and a ‘null’ outcome by activating a third outcome neuron.

Training proceeds in epochs of 128 mini-batches of input-output pairs generated from the first step in the trial (where inputs are the stimulus, action, and outcome in the previous trial, while outputs are the null action and outcome) and the second step in the trial (where inputs are the same as in the first step in addition to the stimulus in the current trial, and the outputs are the correct action and predicted output), randomly intermixed. A fraction of 0.25 inputs-output pairs are generated so as to encode switch trials in the task, i.e. error trials that are following a contextual switch. The implicit assumption here is that the agent does not make mistakes, unless there is a contextual switch, which means that switch trials will be characterized by an outcome indicating an error (see Figure S16a, two rightmost input-output pairs). Errors are encoded with outcome neurons set to a fixed random binary pattern corresponding to a negative outcome in the previous trial, while the target output patterns are chosen so as to correspond to the opposite context compared to the one that was (incorrectly) followed in the previous step.

We train a population of 100 randomly initialized networks on the supervised learning version of the task. Figure S16b shows the resulting learning curves as a function of training epochs in terms of the average accuracy across the population at correctly predicting the null action and outcome after the first step, and correctly predicting the correct action and outcome after the second step, respectively (error bars denote the standard deviation of the accuracy across the population of trained networks).

The main results are analogous to what we observed for the Reinforcement Learning case. Figure S17 reveals that the *context* again cannot be decoded in the input but can already be decoded with high accuracy starting from the randomly initialized first hidden layer. Context becomes abstract in layer L2, where on average CCGP (across the population of models) becomes significantly higher than chance. Correspondingly, the average PS in L2 is fairly high, hovering around 0.25. In both hidden layers of the trained models, the PS and CCGP for context are significantly higher than in the untrained models.

After training, the CCGP and PS for the previous outcome and action variables are very close to the untrained models. However, as we noticed earlier, we note a slight decrease in abstraction of value and action that seems to correlate with an increase in the abstraction of context. We will study this effect in Figure S20.

**Figure S17:**
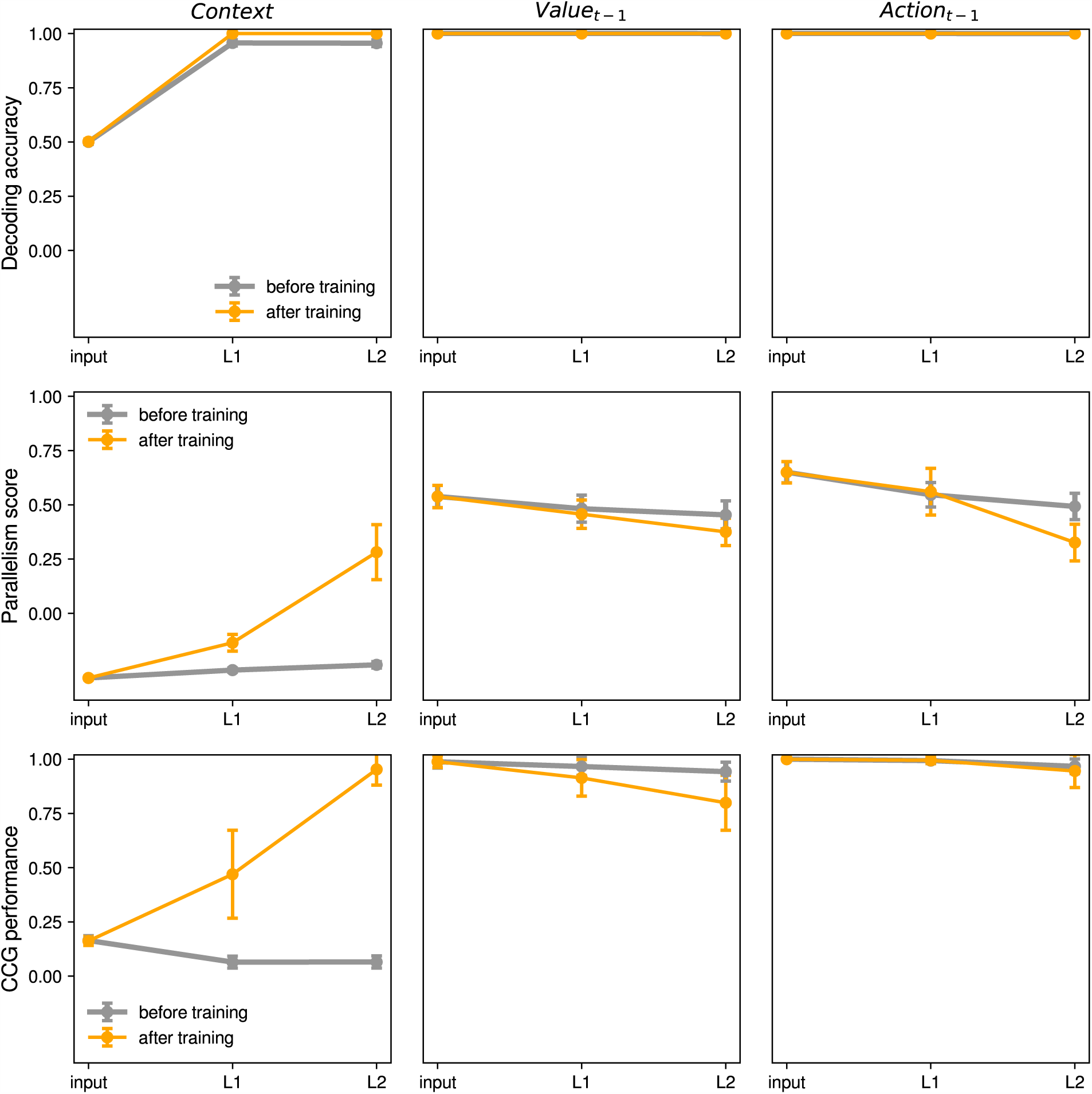
Supervised learning model, activity analysis: *decoding accuracy* (first row), *parallelism score* (second row) and *cross-condition generalization (CCG) performance* (third row) during the simulated first step (pre-stimulus epoch) in the task, for the variables *context* (first column), *previous outcome* (second column) and *previous action* (third column). In each panel we plot one of the quantities of interest as a function of the layer where it is measured in the architecture: inputs, first hidden layer (L1) and second hidden layer (L2). Each point in the plots represents the mean quantity of interest across 100 randomly initialized and independently trained networks, before training (grey data points) and after training (orange data points). Error bars indicate standard deviations computed over the same distribution of 100 networks. The lines connect points computed within the same training condition (before training or after training) in adjacent layers in the architecture.

As in the case of the Reinforcement Learning model, we compare the PS and CCGP of the trained models to the null-distributions obtained from a shuffle of the data and the geometric random model, respectively. Figure S18 shows beeswarm plots of the probability that the CCG performance and parallelism score of each of the trained supervised learning models is above the distributions given by the geometric random model and a shuffle test, respectively, for all variables in the task individually (context, previous value, and previous action) and all variables simultaneously for the same model (All). These probabilities are empirically estimated over 100 trained models and 1000 samples of the null-distributions per trained model. Figure S18 is laid out similarly to Figure S15. As in the Reinforcement Learning case, a considerable fraction of trained models display a PS and CCGP above the 95th percentile of the null-distributions, particularly in hidden layer L2. For instance, in L2 91% of the trained models have parallelism score that is higher than 95% of the shuffle null-distribution simultaneously for all three variables, while 40% have CCG performance above 95% of the null-distribution obtain with the random model.

Next we set out to study the effect that we noticed whereby abstraction of value seems to be negatively correlated with abstraction of context. First, we hypothesize that abstraction of value might be positively correlated with the fraction of switch trials, since maintaining information about the previous outcome (i.e., the value) is crucial to identify a contextual switch. Figure S19 shows scatter plots of the PS and CCGP for value in the hidden layer L2 of 100 supervised learning models trained as in the previous paragraphs but with different fractions of switch trials (between 0 and 0.5), revealing that indeed both quantities are positively correlated with the fraction of switch trials. Next we utilize the fraction of switch trials as an independent variable to control PS and CCGP of value, and quantify the correlation between the observed PS and CCGP for value and context. The first panel of Figure S20 reveals a statistically significant negative correlation −0.34 between the PS for value and the PS for context in hidden layer L2 of the supervised learning model. Analogously, the second panel of Figure S20 shows that the logit transform of the CCGP for context is also negatively correlated with the CCGP for value (we used the logit transform, because CCGP for context tends to saturate close to one for low values of CCGP for value). These results quantify the effect that we had qualitatively alluded to earlier in the Reinforcement Learning model that caused value to be represented in a less abstract format when context is represented in a more abstract format, and which prompted us to introduce the reconstruction loss term.

**Figure S18:**
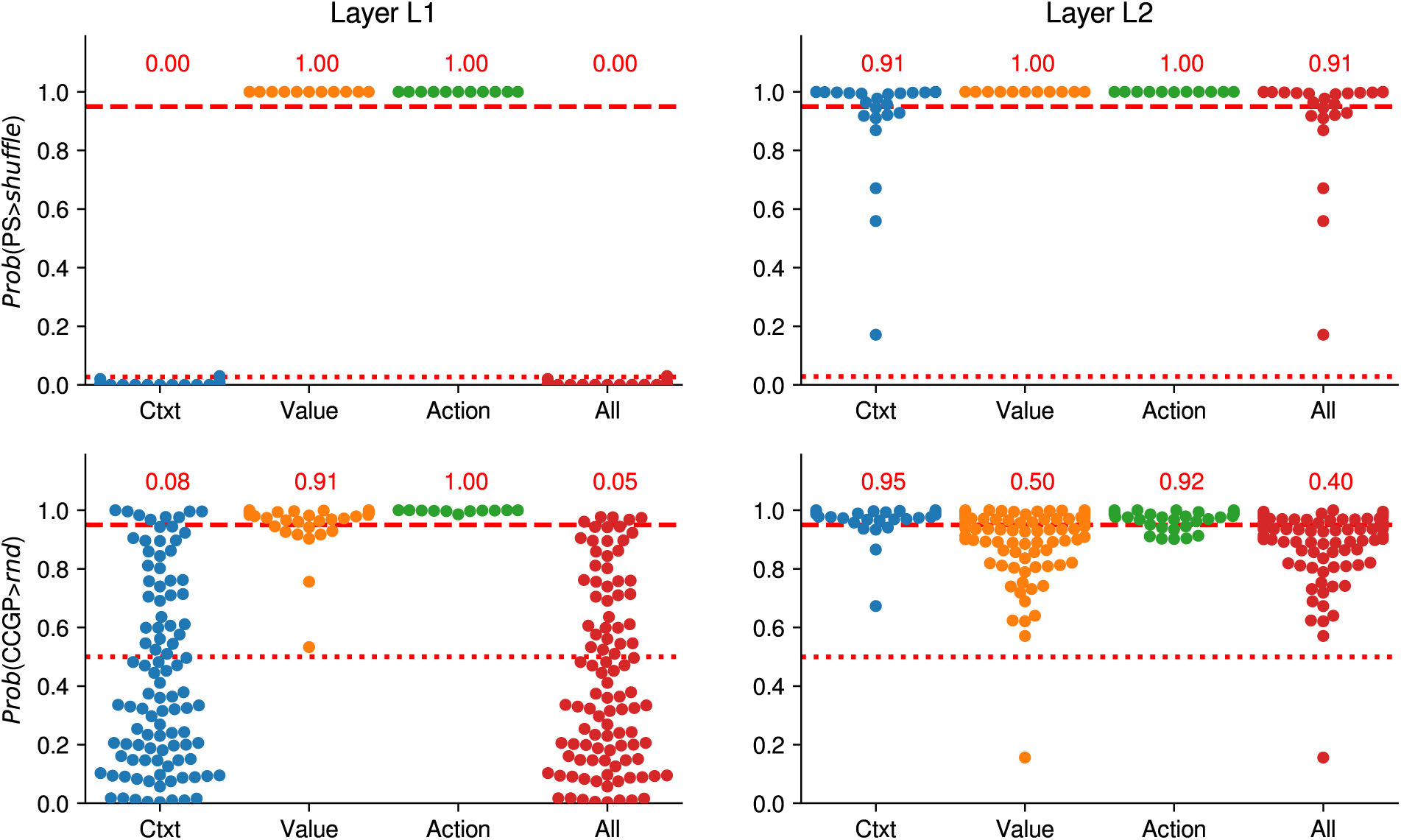
Beeswarm plots of the probability of PS and CCGP of trained supervised learning models to be above the null-distributions given by the geometric random model and a shuffle test of the data (empirically estimated with 1000 samples per model), respectively (see Figure 4). Every dot in the graph represents a trained model. The first row of plots shows the probability that the Parallelism Score of each model for every variable individually (Context, Value and Action) and all variables simultaneously (All) is above the null-distribution given by a shuffle of the data in layer L1 (first column) and L2 (second column). The second row is laid out similarly as the first one, but shows the probability that the CCGP of each model is above the null-distribution given by the geometric random model. The dotted lines corresponds to the mean of the null-distribution, and the dashed line corresponds to the 95th percentile of the null-distribution. The numbers written in red above each plot report the fraction of models that are above the 95th percentile threshold.

**Figure S19:**
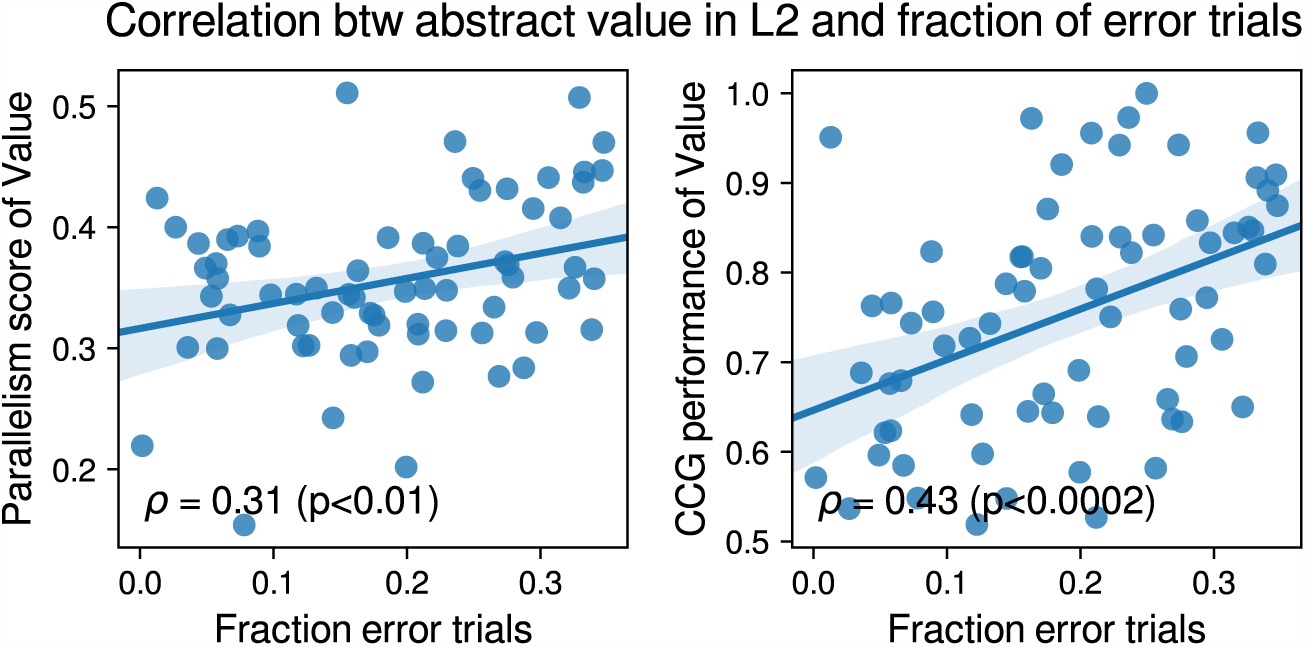
Correlation between abstraction of value and fraction of switch trials in hidden layer L2 of the supervised learning model. Both PS and CCGP for value are positively correlated with the fraction of switch trials with statistically significant Pearson correlation coefficients *ρ* (t-test).

**Figure S20:**
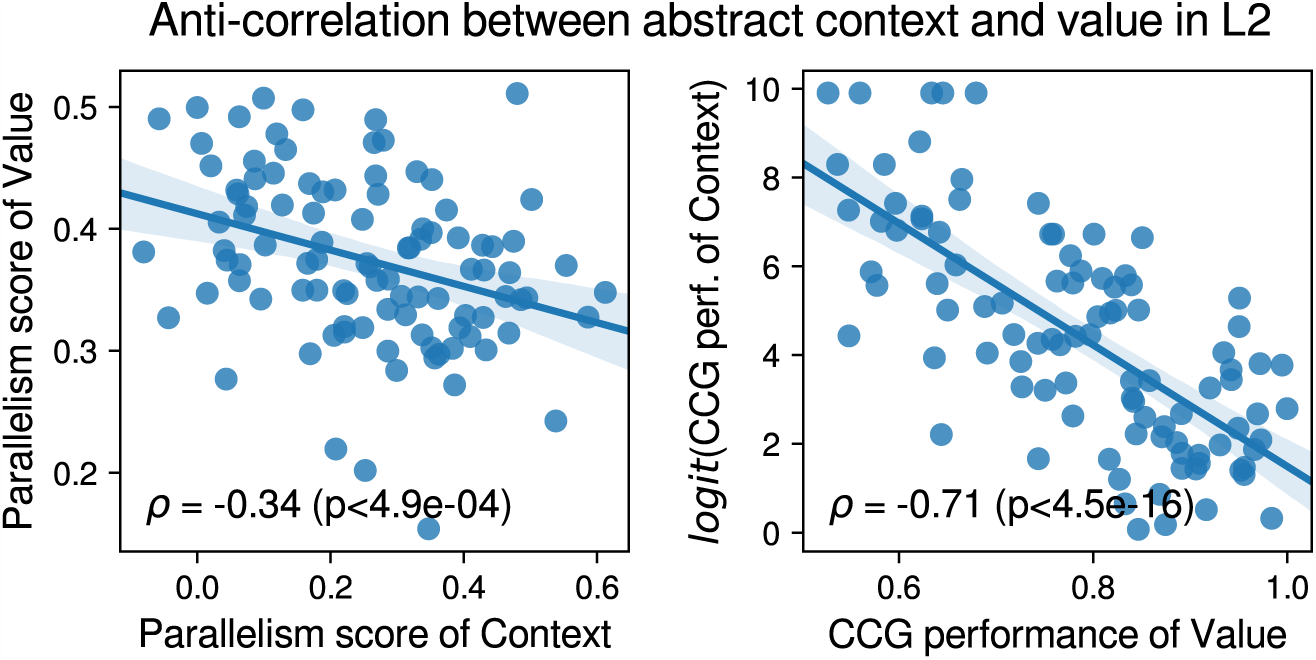
Anti-correlation between abstraction of value and abstraction of context in hidden layer L2 of the supervised learning model. Both PS and CCGP for value are negatively correlated with respectively the PS and CCGP for context. Since CCGP for context tends to saturate close to one for low CCGP for value, we applied a logit transform to the CCGP for context, before computing the Pearson correlation coefficients.

## References

1. Behrens, T. E. et al. What is a cognitive map? organizing knowledge for flexible behavior. Neuron 100, 490–509 (2018).

2. Milner, B., Squire, L. & Kandell, E. Cognitive neuroscience and the study of memory. Neuron 1998, 445–468 (1998).

3. Eichenbaum, H. Hippocampus: Cognitive processes and neural representations that underlie declarative memory. Neuron 2004, 109–120 (2004).

4. Wirth, S. et al. Single neurons in the monkey hippocampus and learning of new associations. Science 300, 1578–1581 (2003).

5. Schapiro, A. C., Turk-Browne, N. B., Norman, K. A. & Botvinick, M. M. Statistical learning of temporal community structure in the hippocampus. Hippocampus 26, 3–8 (2016).

6. Kumaran, D., Summerfield, J., Hassabis, D. & Maguire, E. Tracking the emergence of conceptual knowledge during human decision making. Neuron 63, 889–891 (2009).

7. Wallis, J. D., Anderson, K. C. & Miller, E. K. Single neurons in prefrontal cortex encode abstract rules. Nature 411, 953 (2001).

8. Miller, E. K., Nieder, A., Freedman, D. J. & Wallis, J. D. Neural correlates of categories and concepts. Current opinion in neurobiology 13, 198–203 (2003).

9. Buckley, M. J. et al. Dissociable components of rule-guided behavior depend on distinct medial and prefrontal regions.. Science 325, 52–58 (2009).

10. Antzoulatos, E. G. & Miller, E. K. Differences between neural activity in prefrontal cortex and striatum during learning of novel abstract categories. Neuron 71, 243–249 (2011).

11. Wutz, A., Loonis, R., Roy, J. E., Donoghue, J. A. & Miller, E. K. Different levels of category abstraction by different dynamics in different prefrontal areas. Neuron 97, 716–726 (2018).

12. Saez, A., Rigotti, M., Ostojic, S., Fusi, S. & Salzman, C. Abstract context representations in primate amygdala and prefrontal cortex. Neuron 87, 869–881 (2015).

13. Rigotti, M. et al. The importance of mixed selectivity in complex cognitive tasks. Nature 497, 585 (2013).

14. Fusi, S., Miller, E. K. & Rigotti, M. Why neurons mix: high dimensionality for higher cognition. Current opinion in neurobiology 37, 66–74 (2016).

15. Mnih, V. et al. Human-level control through deep reinforcement learning. Nature 518, 529 (2015).

16. Yakovlev, V., Fusi, S., Berman, E. & Zohary, E. Inter-trial neuronal activity in inferior temporal cortex: a putative vehicle to generate long-term visual associations. Nature neuroscience 1, 310 (1998).

17. Bernacchia, A., Seo, H., Lee, D. & Wang, X.-J. A reservoir of time constants for memory traces in cortical neurons. Nature neuroscience 14, 366 (2011).

18. Akrami, A., Kopec, C. D., Diamond, M. E. & Brody, C. D. Posterior parietal cortex represents sensory history and mediates its effects on behaviour. Nature 554, 368 (2018).

19. Stringer, C., Pachitariu, M., Steinmetz, N., Carandini, M. & Harris, K. D. High-dimensional geometry of population responses in visual cortex. bioRxiv 374090 (2018).

20. Gao, P. & Ganguli, S. On simplicity and complexity in the brave new world of large-scale neuroscience. Current opinion in neurobiology 32, 148–155 (2015).

21. Mazzucato, L., Fontanini, A. & La Camera, G. Stimuli reduce the dimensionality of cortical activity. Frontiers in systems neuroscience 10, 11 (2016).

22. Tang, E., Mattar, M. G., Giusti, C., Thompson-Schill, S. L. & Bassett, D. S. Effective learning is accompanied by increasingly efficient dimensionality of whole-brain responses. arXiv preprint 1709.10045 (2017).

23. Lindsay, G. W., Rigotti, M., Warden, M. R., Miller, E. K. & Fusi, S. Hebbian learning in a random network captures selectivity properties of the prefrontal cortex. The Journal of neuroscience: the official journal of the Society for Neuroscience 37, 11021–11036 (2017).

24. Bellman, R. E. Dynamic Programming. (Princeton University Press, 1957).

25. Dietterich, T. G. Hierarchical reinforcement learning with the maxq value function decomposition. Journal of Artificial Intelligence Research 13, 227–303 (2000).

26. Precup, D. Temporal abstraction in reinforcement learning (PhD thesis, University of Massachusetts Amherst, 2000).

27. Barto, A. G. & Mahadevan, S. Recent advances in hierarchical reinforcement learning. Discrete Event Dynamic Systems 13, 341–379 (2003).

28. Ponsen, M., Taylor, M. E. & Tuyls, K. Abstraction and generalization in reinforcement learning: A summary and framework. In International Workshop on Adaptive and Learning Agents, 1–32 (Springer, 2009).

29. Mikolov, T., Yih, W.-t. & Zweig, G. Linguistic regularities in continuous space word representations. In Proceedings of the 2013 Conference of the North American Chapter of the Association for Computational Linguistics: Human Language Technologies, 746–751 (2013).

30. Mikolov, T., Sutskever, I., Chen, K., Corrado, G. S. & Dean, J. Distributed representations of words and phrases and their compositionality. In Advances in neural information processing systems, 3111–3119 (2013).

31. Mitchell, T. M. et al. Predicting human brain activity associated with the meanings of nouns. science 320, 1191–1195 (2008).

32. Chen, X. et al. Infogan: Interpretable representation learning by information maximizing generative adversarial nets. In Advances in neural information processing systems, 2172–2180 (2016).

33. Higgins, I. et al. β-VAE: Learning basic visual concepts with a constrained variational framework. In ICLR (2017).

34. Chen, T. Q., Li, X., Grosse, R. B. & Duvenaud, D. K. Isolating sources of disentanglement in variational autoencoders. In Advances in Neural Information Processing Systems 31, 2614–2624 (2018).

35. Kim, H. & Mnih, A. Disentangling by factorising. arXiv preprint 1802.05983 (2018).

36. Riesenhuber, M. & Poggio, T. Hierarchical models of object recognition in cortex. Nature neuroscience 2, 1019 (1999).

37. LeCun, Y., Bengio, J. & Hinton, G. Deep learning. Nature 521, 436–444 (2015).

38. Freedman, D. J., Riesenhuber, M., Poggio, T. & Miller, E. K. Categorical representation of visual stimuli in the primate prefrontal cortex. Science 291, 312–316 (2001).

39. Rust, N. & Dicarlo, J. Selectivity and tolerance (“invariance”) both increase as visual information propagates from cortical area v4 to it. J Neurosci 30, 12978–12995 (2010).

40. DiCarlo, J. J. & Cox, D. D. Untangling invariant object recognition. Trends in cognitive sciences 11, 333–341 (2007).

41. DiCarlo, J. J., Zoccolan, D. & Rust, N. C. How does the brain solve visual object recognition? Neuron 73, 415–434 (2012).

42. Isik, L., Meyers, E. M., Leibo, J. Z. & Poggio, T. The dynamics of invariant object recognition in the human visual system. Journal of neurophysiology 111, 91–102 (2013).

43. Isik, L., Tacchetti, A. & Poggio, T. A fast, invariant representation for human action in the visual system. Journal of neurophysiology 119, 631–640 (2017).

44. Meyers, E. M., Freedman, D. J., Kreiman, G., Miller, E. K. & Poggio, T. Dynamic population coding of category information in inferior temporal and prefrontal cortex. Journal of neurophysiology 100, 1407 (2008).

45. Golland, P., Liang, F., Mukherjee, S. & Panchenko, D. Permutation tests for classification. In International Conference on Computational Learning Theory, 501–515 (Springer, 2005).

46. Borg, I. & Groenen, P. Modern multidimensional scaling: Theory and applications. Journal of Educational Measurement 40, 277–280 (2003).

47. Stefanini, F. et al. A distributed neural code in ensembles of dentate gyrus granule cells. bioRxiv 292953 (2018).

48. Morcos, A. S., Barrett, D. G., Rabinowitz, N. C. & Botvinick, M. On the importance of single directions for generalization. arXiv preprint 1803.06959 (2018).

49. Machens, C. K., Romo, R. & Brody, C. D. Functional, but not anatomical, separation of “what” and “when” in prefrontal cortex. J Neurosci 30, 350–360 (2010). URL http://dx.doi.org/10.1523/JNEUROSCI.3276-09.2010.

50. Chen, X. Confidence interval for the mean of a bounded random variable and its applications in point estimation. arXiv preprint 0802.3458 (2008).

51. Kingma, D. P. & Ba, J. Adam: A method for stochastic optimization. arXiv preprint 1412.6980 (2014).

52. Paszke, A. et al. Automatic differentiation in pytorch. In NIPS 2017 Autodiff Workshop (2017).

53. Rigotti, M., Ben Dayan Rubin, D., Wang, X.-J. & Fusi, S. Internal representation of task rules by recurrent dynamics: the importance of the diversity of neural responses. Frontiers in Computational Neuroscience 4, 24 (2010).

54. Barak, O., Rigotti, M. & Fusi, S. The sparseness of mixed selectivity neurons controls the generalization-discrimination trade-off. The Journal of neuroscience: the official journal of the Society for Neuroscience 33, 3844–3856 (2013).

